# A single layer artificial neural network type architecture with molecular engineered bacteria for complex conventional and reversible computing

**DOI:** 10.1101/2021.08.05.455238

**Authors:** Kathakali Sarkar, Deepro Bonnerjee, Rajkamal Srivastava, Sangram Bagh

## Abstract

Here, we adapted the basic concept of artificial neural networks (ANN) and experimentally demonstrate a broadly applicable single layer ANN type architecture with molecular engineered bacteria to perform complex irreversible computing like multiplexing, de-multiplexing, encoding, decoding, majority functions, and reversible computing like Feynman and Fredkin gates. The encoder and majority functions and reversible computing were experimentally implemented within living cells for the first time. We created molecular-devices, which worked as artificial neuro-synapses in bacteria, where input chemical signals were linearly combined and processed through a non-linear activation function to produce fluorescent protein outputs. To create such molecular devices, we established a set of rules by corelating truth tables, mathematical equations of ANN, and molecular-device design, which unlike molecular computing, does not require circuit diagram and the equation directly correlates the design of the molecular-device. To our knowledge this is the first adaptation of ANN type architecture with engineered cells. This work may have significance in new platform for biomolecular computing, reversible computing and in transforming living cells as ANN-enabled hardware.

## Introduction

Hardware implementation of artificial neural network (ANN) through neuro-synapse type architectures^1–4^ have been developed and it has important implications in creating intelligent autonomous systems^5,6^. However, ANN hardware is mainly limited to various neuromorphic chips made out of various inorganic materials^1–10^, photonics^8^, spintronics^9^, and in-vitro DNA computation^10,11^. The physical mechanism of ANN hardware operation is strikingly simpler than the biological neurons^12^ and their networks in the brain^13^. Though, our understanding of neural networks in the brain is far from complete^13^, our understanding of ANN is from the first principle^12^. This may allow adaptation of the basic ANN type architecture with living cells by engineering interactions at the molecular level.

In this study, we experimentally created a broadly applicable single layer ANN type framework using engineered molecular devices in living bacteria for performing complex conventional irreversible and nonconventional reversible computation. Here, the molecular devices inside bacteria work as artificial neuro-synapses and we named it as ‘bactoneuron’ (BNeu). The molecular devices, linearly combine the chemical inputs and transforms it nonlinearly to a fluorescent protein output. We experimentally demonstrated that single-layer neural network type architectures stemmed from those bactoneurons were general, flexible and perform complex irreversible (conventional) computation through a 2-to-4 decoder, a 4-to-2-priority encoder, a majority function, a 1-to-2 de-multiplexer, and a 2-to-1 multiplexer^14,15^ and reversible logic mapping through Feynman and Fredkin Gate. To our knowledge, the encoder and majority function have not been demonstrated and reversible computing is never explored in living biological systems. We established a set of molecular principles by corelating truth tables, mathematical equations of ANN, and molecular-device design, such that unlike invitro and in-vivo molecular computation, our ANN framework showed a complementary way to design biomolecular computation without following the electronic hierarchical logic circuit design principles and a single mathematical equation based on its sign of the parameters directed the design of the molecular device.

## Results and Discussions

### Principles of mapping functional truth table to single layer molecular engineered bacterial ANN

First, we hypothesized that an abstract ANN model can be mapped into an engineered cellular model (Fig.1a), where engineered molecular devices inside bacterial cell work as artificial neuro-synapse (bactoneurons). The bactoneurons combined the inputs in the form of environmental chemical inputs and executed appropriate log-sigmoid activation functions (equation 1) through engineered molecular devices. Equation 1 is a conventional activation function for characterization of an artificial neuro-synapse in ANN^12^ with two inputs and can be applied to a wide range of functional behaviors, based on its sign and magnitudes of the weight and bias terms.

**Figure 1:**
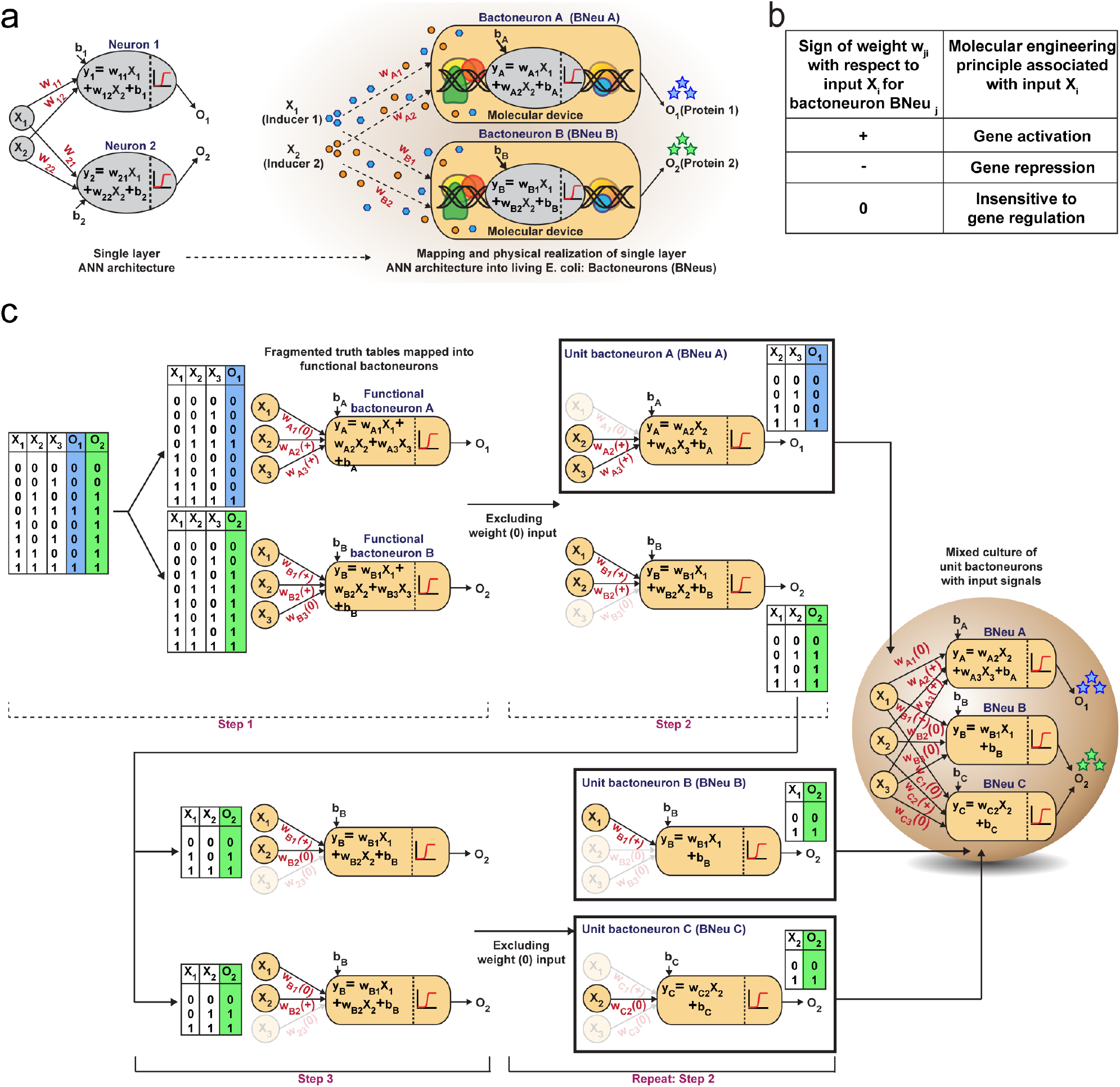
Single layer artificial neural network (ANN) type architechture with engineered bacteria and schematic design rules of building complex computing function. **a)** Schematic representation of an ideal single layer ANN with two weighted inputs (X_1_ and X_2_) and outputs (O_1_ and O_2_) along with their corresponding weights (w_i_), the biases (b_i_) and summation function (y_i_). This abstract ANN is mapped with the proposed artificial bacterial neurons (Bactoneurons or BNeus). **b)** Relation between signs of weights in the activation function and molecular engineering principles for a bactoneuron associated with a input. **c)** Making of bacteria-based single layer ANN type architecture from truth table of a given function.

In this equation, each bactoneuron had two weight values of varying signs and magnitude corresponding to its two chemical inducer inputs (X_1_ and X_2_) and a bias with its value in accordance with the functional response of the neuron. For a given bactoneuron j,

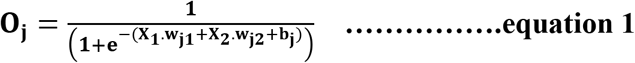

where,

O_j_ is the output from neuron j,

X_1_ and X_2_ represent two input inducer concentration,

W_j1_ represents the weight of input X_1_ for the neuron j,

W_j2_ represents the weight of input X_2_ for the neuron j,

b_j_ represents the bias for the neuron j.

Equation 1 suggests that if w_J1_ is positive, output O_i_ would increase with X_1_. In terms of molecular device, it is similar to a molecular activation (Fig.1b). Similarly, negative weight would suggest a repression and ‘zero’ weight suggests the insensitivity of the input with the output (Fig.1b). This simple correlation between the sign of a weight, w_ji_ within an activation function of a ‘unit’ bactoneuron and molecular engineering principle, guided the physical design of the molecular device.

Next, we devised a way to map a complex computing function through a single layer ANN type architecture directly from its functional truth table, without considering its hierarchical electronic design principle (Fig.1c). We built a set of rules to derive bactoneurons from the functional truth table by dividing the bigger truth tables into smaller ones and to connect them with bactoneuron design (Fig.1c). We demonstrated this process considering a random functional truth table (Fig.1c). First, we considered a single output within a functional truth table and then looked at its relation with all the input combinations. We grouped those input combinations in the form of a smaller truth tables, in such a way, that each input corresponding to that particular output possessed a weight with only one type of sign (+, − or 0) (step 1). Such individual bactoneurons in this step were named as ‘functional bactoneurons’, which in appropriate ANN combinations would give rise to the actual function. Next, we ignored the weight(s) with ‘zero’ values, if any, from functional bactoneurons and mapped them back with smaller truth tables (step 2). Further, we looked at the output of the smaller truth table from step2. If the output value 1 (true) appeared only once in the smaller truth tables, we defined them as ‘unit bactoneurons’. Otherwise, we kept dividing the truth table (step 3), until the above condition appeared. This way we identified the unit bactoneurons, which is the smallest unit required to be constructed as molecular device. Once those unit bactoneurons are combined appropriately, they would operate as the functional bactoneurons. Thus, when the unit bactoneurons were assembled according to ANN structure, the actual function was physically realized (Fig.1c).

Now, we chose a range of computing functions with varying complexities (Fig.2) and derived their functional bactoneurons (Fig.2) from their functional truth table following the principle stated above (Fig.1c). The chosen functions included 1-to-2 demultiplexer^14^ (Fig.2a), 2-to-1 multiplexer^14^ (Fig.2b), majority functions^15^ (Fig.2c), 2-to-4 decoder^14^ (Fig.2d), and 4-to-2 priority encoder^14^ (Fig.2e). A de-multiplexer performs as an output selector where it takes input from just one source and the logical state(s) of selector line(s) direct(s) to select only one among multiple output channels to process the signal and interpret it. Multiplexer performs the complementary function where the logical state(s) of selector line(s) directs which input is to be received for generating an output. A majority function suggests that in a ternary system if more than 50% of the inputs are true then the output is true, otherwise false. A N:2N decoder converts N bit binary-coded inputs into 2N coded outputs in a one-to one mapping fashion and an encoder encodes input signals to fewer bits and transforms them into encoded outputs. The details of deriving functional and unit bactoneurons from the truth tables of all those functions without considering its integrated circuit design are shown in figure 2 and supplementary figure S1. The specific activation function equations for all functional bactoneurons are shown in table S1.

**Figure 2:**
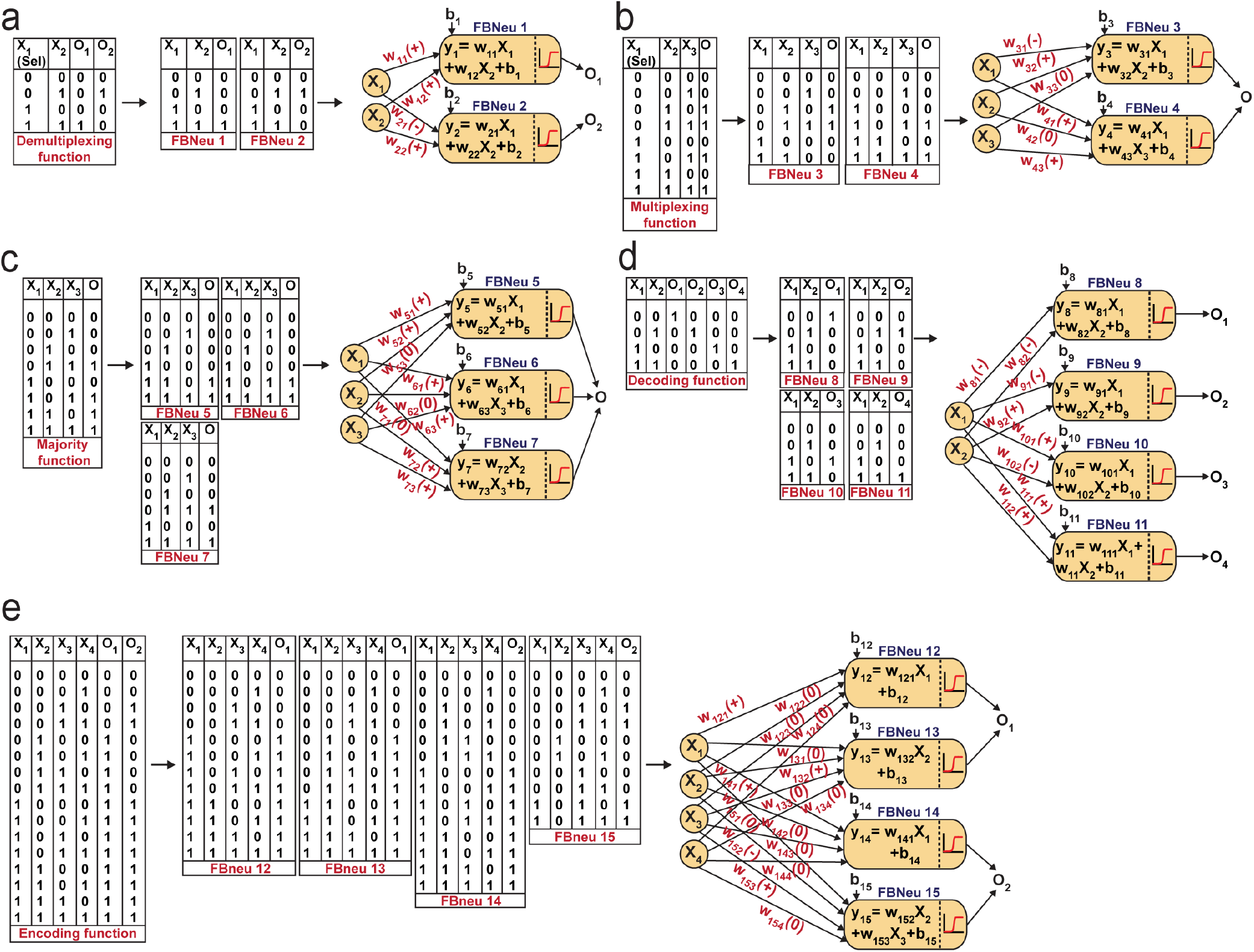
Derivation of functional bactoneurons and abstraction of signle layer ANN type arcitechtures from functional truth tables of complex computing functions. **a)** 1-to-2 de-multiplexer, **b)** 2-to-1 multiplexer, **c)** 3-input majority function, **d)** 2-to-4 decoder and **e)** 4-to-2 priproty encoder.

### Design and construction of molecular devices and unit bactoneurons

Next, we developed molecular devices to construct the unit bactoneurons (Table-S1) in our chassis organism, E. coli DH5αZ1^16^. The molecular devices are engineered molecular networks incorporated in plasmid vectors, which are 25 nm circular DNAs replicate inside the bacteria^17^. In our design, the abstract inputs (X_i_) were replaced by extracellular chemical inducers like Isopropyl β-D-1 thiogalactopyranoside (IPTG), anhydrotetracycline (aTc), N-Acyl homoserine lactone (AHL), and arabinose. The abstract outputs (O_i_) were changed to fluorescent proteins like EGFP, mKO2, E2 Crimson, mTFP1, and mVenus as appropriate. The device design of the unit bactoneurons were based on the molecular engineering principle we stated in figure 1b.

We started with the construction and characterization of unit bactoneuron BNeu 1 (Fig.3a-d), where both the weights in the activation function are positive with respect to the inputs (X_1_ and X_2_). In BNeu 1, two inducer chemicals IPTG and aTc were used as the inputs X_1_ and X_2_ respectively while enhanced green fluorescence protein (EGFP) was used as the output O_1_. The molecular device (Fig.3a) for BNeu 1 consists of a synthetic hybrid promoter, which combines the chemical signals aTc and IPTG and processes them through a log-sigmoid function (equation 1) and according to the principle, both aTc and IPTG should work as activator for the system. TetR and LacI, two transcription factors, which are constitutively and endogenously expressed in *E. coli* DH5αZ1, bind the hybrid promoter thereby hindering it from expressing EGFP^16^. Both IPTG and aTc, which bind with LacI and TetR respectively and change their conformation such that they cannot bind to the promoter anymore, the promoter is free to recruit RNA polymerase for EGFP expression.

**Figure 3:**
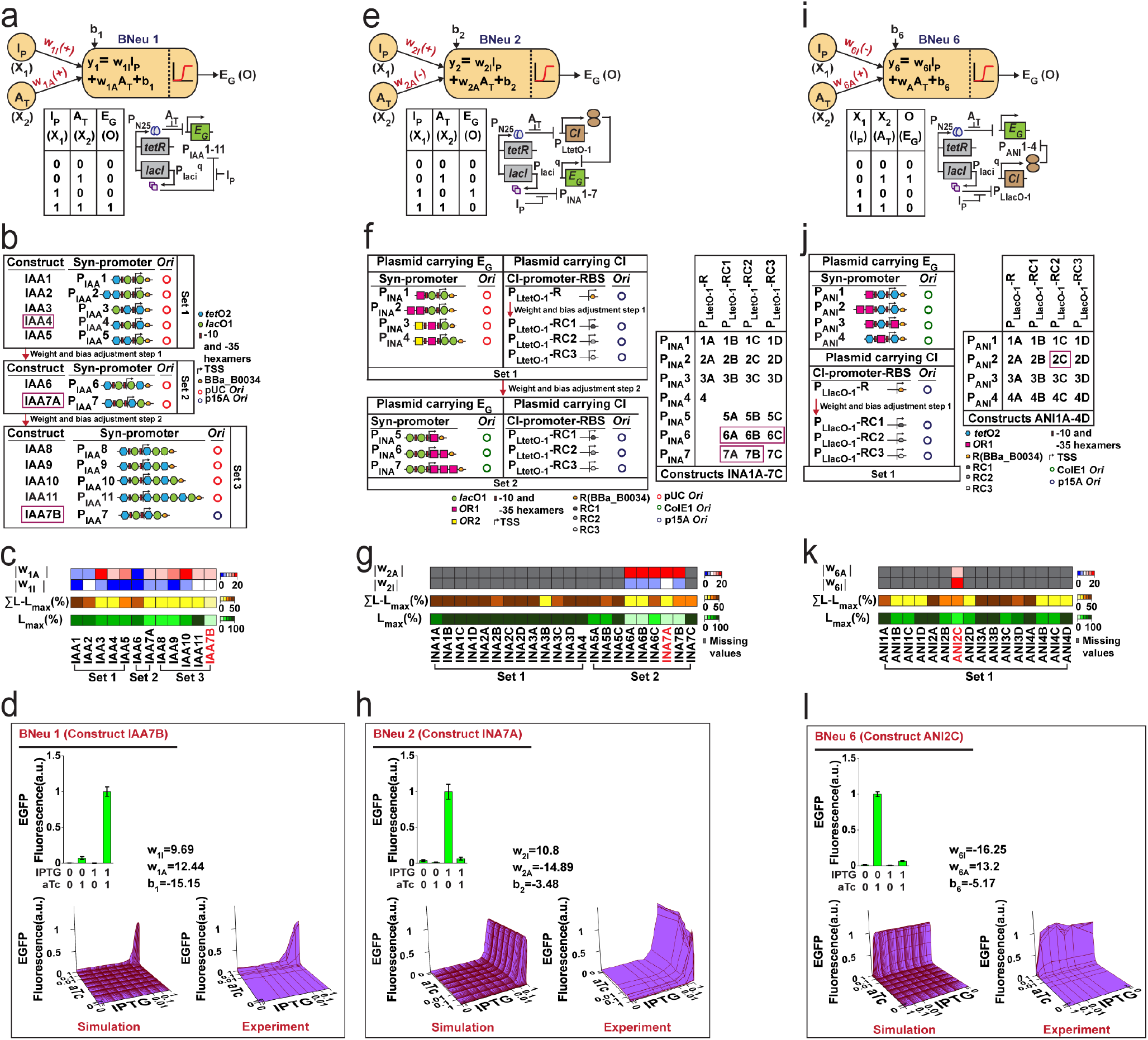
Design, experimental characterization, and weight and bias adjustems of molecular devices for unit bactoneurons BNeu 1, BNeu 2 and BNeu 6. **a)** Neural architecture of the bactoneuron BNeu 1 where w_1I_ and w_1A_ are weights of inputs IPTG (I_P_: X_1_) and aTc (A_T_: X_2_) and b_1_is the bias. Activation function generates output EGFP (E_G_: O). Truth table and biological circuit design of the BNeu 1 activation function. P_IAA_1-11 are synthetic promoters designed for BNeu 1 activation function and regulated by IPTG and aTc as these promoters contain binding cites for LacI and TetR proteins. **b)** Schematic representation of weight and bias adjusment of BNeu 1 through constructs IAA1-11. IAA1-11 were generated in sets by changing hybrid promoter design with varied number and positions of the operating sites for both TetR and LacI and by simultaneously changing the origin of replication (Ori) of the plasmids carrying those promoters. Promoter maps of P_IAA_1-11 are also shown. Positions of −10 and −35 hexamers, transcription start site, ribosome binding sites (RBS) and LacI & TetR binding sites are depicted in individual promoter maps. Weights and bias associated with IPTG and aTc of the BNeu 1 were adjusted through a two-step modification of molecular interactions. Constructs corresponding to the selected promoter from each set are shown in magenta box. **c)** Heatmaps showing percentage highest leakage (L_max_(%)), percentage sum of leakage excluding highest leakage (ΣL-L_max_(%)), modulus of weight associated with IPTG (|w_1I_|) and modulus of weight associated with aTc (|w_1A_|) for each out of 12 constructs (constructs IAA1-11). Construct IAA7B (coloured in red) is selected as the best performer. **d)** Expression charectarization, simulated behavior (3D plot), and experimental validation of unit bactoneuron BNeu 1 carrying construct IAA7B are shown. All of these experimental data were collected after 10h induction follwed by resuspension and 6h induction. **e)** Neural architecture, truth table and biological circuit design of the bactoneuron BNeu 2 activation function. **f)** Weight and bias adjustment of BNeu 2 through constructs INA1A-7C, which are a two-plasmid system. Here, we re-engineered the synthetic promoter of BNeu 1 and replaced the aTc gene-activation function (+w) with aTc gene repression function (−w), such that in the presence of aTc, transcription from the promoter gets turned off. In this promoter we introduced an operating site for λ repressor CI proteins and the amount of CI was under the control of an aTc-inducible promoter. Promoter maps, *Ori,* RBS, and its various combinations in constructs are shown. **g)** Heat maps of weight, leakage and hence bias adjustment for BNeu 2. **h)** Characterization, simulation and validation of BNeu 2 (INA1A-7C). **i)** Neural architechture, truth table and biological design of BNeu 6. **j)** Details of the constructs (ANI1A-4D) for weight and bias adjustment of BNeu 6 and **k)** corresponsing heat maps are also shown. Construct ANI2C was chosen as best performing construct. **l)** Characterization, simulation and validation of BNeu 6 construct ANI2C.

To start with random weights and biases, as in any ANN design^12^, we constructed and characterized an initial set (Set 1) of molecular constructs (IAA1-5), (Fig. 3b, Table-S2) containing synthetic promoters P_IAA_1-P_IAA_5 respectively (Table-S3). For this, we measured the EGFP expression at various combinations of ‘zero’ and ‘saturated’ concentration of IPTG and aTc (Fig.S2a) and performed dose responses (Fig.S2b) of EGFP expression as a function of IPTG/aTc, by varying the concentration of one chemical, while keeping the other at saturated concentration. The dose-response behaviors were fitted to a modified form of equation 1 (equation-2, supplementary note-1). The fitting parameters gave the values of weights for each input and biases (Table S4). This starting set (Set 1) showed high leakage and lower weight values (Fig. 3c, Fig. S2a, Fig. S2b, table-S4 and table-S5).

Higher values for w_1I_, w_1A_ would signify sharper transition from OFF to ON state and more negative bias might signify reduced intercept in a dose-response curve, which could get reflected as less ‘leakage’, which was defined by the basal EGFP expression.

Guided by the least leakage and reasonably high weight values for both w_1I_ and w_1A_, we chose construct IAA4 where promoter P_IAA_4 served as a platform and adjusted the weights and bias by tweaking the molecular design and further creating new constructs (Fig.3b, Table-S2, S3). We iteratively performed this adjustment process a few times (Fig.3c). Clearly, the weight ‘w’ of a bactoneuron was a strong function of types and degree of molecular interactions, as evident from the fact that the weights of the initial bactoneuron were adjusted to a new one in each iteration. We found w_1I_ as the limiting weight as this had a lower value than w_1A_ (Fig.S2b, table-S4). In addition, we focused on the highest leakage value (L_max_), and (ΣL-L_max_) of each construct, where ΣL is the total leakage (Table-S5). Our goal was to reduce it. Constructs IAA4 and IAA5 from the first set carried similar weight but IAA5 showed significantly high ΣL-L_max_. Thus, IAA4 was chosen for further adjustment. Figure 3c shows the adjustment of values for weights and leakage. The constructs for further adjustment in each step are boxed. Now, the construct IAA7A (for BNeu 1) from the second set of adjustment, was taken for further weight adjustment either by engineering the promoter or by altering the relative numbers of the promoters per cell by changing the copy number of the plasmids (Fig.3b). Construct IAA7B (Table-S2) had higher weight values and least leakage with respect to construct IAA7A (Fig. 3c, table-S3 and table-S4). Others (IAA8-11) from the same iteration showed comparatively poor behavior. Such scenario could be compared with overshooting of weight adjustment, as happens in ANN^12^. Thus, molecular device IAA7B was selected as the unit bactoneuron BNeu 1. We performed a simulation and experimentally tested the behavior of the BNeu 1 by simultaneously changing the concentration of the IPTG and aTc (Fig.3d). The results show a close topological match with the simulation (Fig.3d).

In an ANN framework, bias may determine the intercept^12^. The leakage in bactoneuron determines the intercept in the dose-response curves. A moderate correlation (R^2^=0.76) between the bias ‘b_1_’ and L_max_ was found (Fig. S3a) within the experimental range for BNeu 1 construct IAA7B. However, in this case, adjustments of weight values were linked to that of the bias during iteration and was difficult to distinguish the exact molecular reasoning. We performed a simulation by varying the bias but keeping the weights constant and it suggested that, ‘bias’ manifested as leakage in BNeu 1 (Fig.S3b). We performed similar simulations (Fig. S3) for all the unit bactoneurons (Table-S1) and the results suggested that the bias value in bactoneuron indicated the leakage from the molecular devices within a parameter range.

We further illustrate the construction of two other bactoneurons namely BNeu 2 and BNeu 6, which showed positive weight for one inducer and negative for the other (Fig.3e-h, and 3i-l). The development of BNeu 1 indicated that, L_max_ could be the first parameter to look for during molecular engineering. Therefore, for BNeu 2 and BNeu 6, we first checked if the fold change between the highest signal (output logic level “1”) and the highest leakage was more than an arbitrarily threshold 8 times (Table S5). If it was so, we proceeded to adjust the weight values and linked-biases for optimal behavior of the corresponding unit bactoneurons either by engineering the cis-trans element interaction on the promoter, or by reducing the translation rate of CI via RBS designing (Table-S6) or by modulating the relative number of synthetic promoters per cell through modification of the plasmid copy number. We followed this processing pipeline to characterize, fit and adjust the weights and biases in iterations to get the unit bactoneurons BNeu 2 and BNeu 6 (Fig.3e-h and Fig.3i-l), executed by constructs INA7A and ANI2C respectively (Table-S2). The gene expression characterizations, dose-response and fitting for all constructs for BNeu 2 and BNeu 6 were shown in figure S2c,d and S2e,f respectively. The design, gene expression characterizations, dose-response, fitting, simulation, and experimental validation of all other unit bactoneurons are shown in the figure S4. In many cases the unit bactoneurons were equivalent to the functional bactoneurons which did not have any ‘0’ weight inducer input (insensitive to a certain inducer input). Therefore, for such bactoneurons, we experimentally validated the weight ‘zero’ characteristics with respect to the appropriate inputs (Fig.S5).

### ANN created from molecular engineered bactoneurons generate complex computing function

For unit bactoneuron construction, we used EGFP as an output. We changed the EGFP with mKO2, E2-Crimson, mTFP1, and mVenus (Table-S1) as appropriate. Next, the unit bactoneurons were mixed, cocultured and exposed to various combinations of input chemicals following relevant ANN designs (Fig.2). The experimental results are shown in figure 4 and figure S6 for 1-to-2 de-multiplexer (Fig.4a,b, Fig.S6a), 2-to-4 multiplexer (Fig.4c,d, Fig.S6 b), 3-input majority function (Fig.4e,f, Fig.S6c), 2-to-4 decoder (Fig.4g,h, Fig.S6 d), and 4-to-2 priority encoder (Fig.4i,j, Fig.S6e). The results showed the expected truth table behavior (Fig.4).

**Figure 4:**
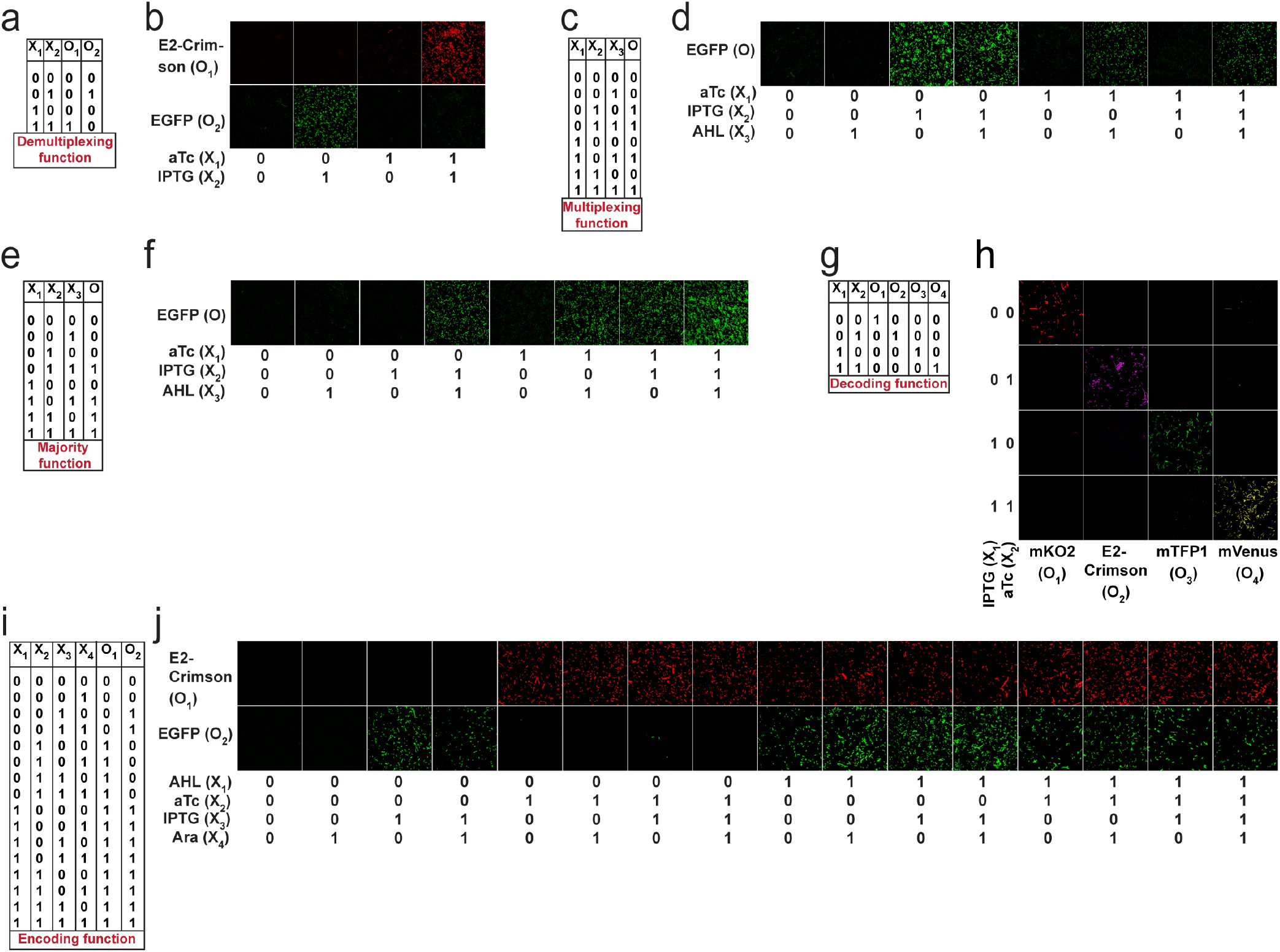
Experimental demonstration of complex computations wirh single layer ANN type architectures with molecular engineered bactoneurons. Truth tables of the **a)** 1-to-2 de-multiplexer, **c)** 2 to-1 multiplexer, **e)** 3-input majority function, **g)** 2-to-4 decoder and **i)** 4-to-2 priority encoder are shown. Inputs and outputs are written as X_i_ (i=1 to n) and O_i_ (i=1 to n) respectively. Experimental behavior of the bacteria-based single layer ANN type architectures corresponding to the **b)** 1-to-2 de multiplexer, **d)** 2-to-1 multiplexer, 3-input majority function, **h)** 2-to-4 decoder and **j)** 4-to-2 priority encoder, studied with fluorescence microscope. Unit bactoneuorns of a function were cultured in a mixed population and treated with all possible combinations of inputs. The resultant expression of fluorescent proteins are represented by separate output channels.

### Mapping logical reversibility with molecular engineered bactoneurons: Feynman and Fredkin Gates

Reversible computing is the heart of quantum computing^18^ and it can map the previous state of the computation from the current state in a one-to-one basis^18,19^. This is called logical reversibility, which was demonstrated in by implementing logically reversible Fredkin and Toffoli gates through in-vitro DNA computation^20^. However, no logically reversible gate is implemented in living cells. Though, the thermodynamic reversibility of reversible computing, which gives lowest energy cost in computation, is not possible in living systems, the potential of logical reversibility in biological systems is yet to explore. We showed that ANN with bactoneurons had the flexibility to create reversible computing and we demonstrated the universal reversible Feynman gate (Fig.5a-b, Fig.S1f, S6f) and Fredkin gate (Fig.5c-d, Fig.S1g, S6g), which may create any linear reversible logic gate. First we derived the functional and unit bactoneurons for Feynman (Fig.5a, fig.S1f, table-S1) and Fredkin gate (Fig.5c, Fig.S1g, table-S1). Feynman gate was represented by 3 unit bactoneurons (BNeu2, BNeu6, BNeu8), which we already developed. The ANN created from the corresponding functional bactoneurons (FBNeus 16-18) of these unit bactoneurons (Fig.5a) showed successful Feynman gate (Fig.5b). Fredkin gate was represented by 5 unit bactoneurons (BNeu 3, 4, 7, 9, 10), where BNeu 3, 4, 7 were already developed and BNeu 9 and BNeu 10 were created (Fig.S4p-u). The ‘zero’ weights of the bactoneurons, where appropriate, was validated in figure S5. The ANN, created from the corresponding functional bactoneurons (FBNeus 19-23) (Fig.5c), showed successful Fredkin gate (Fig.5d).

**Figure 5:**
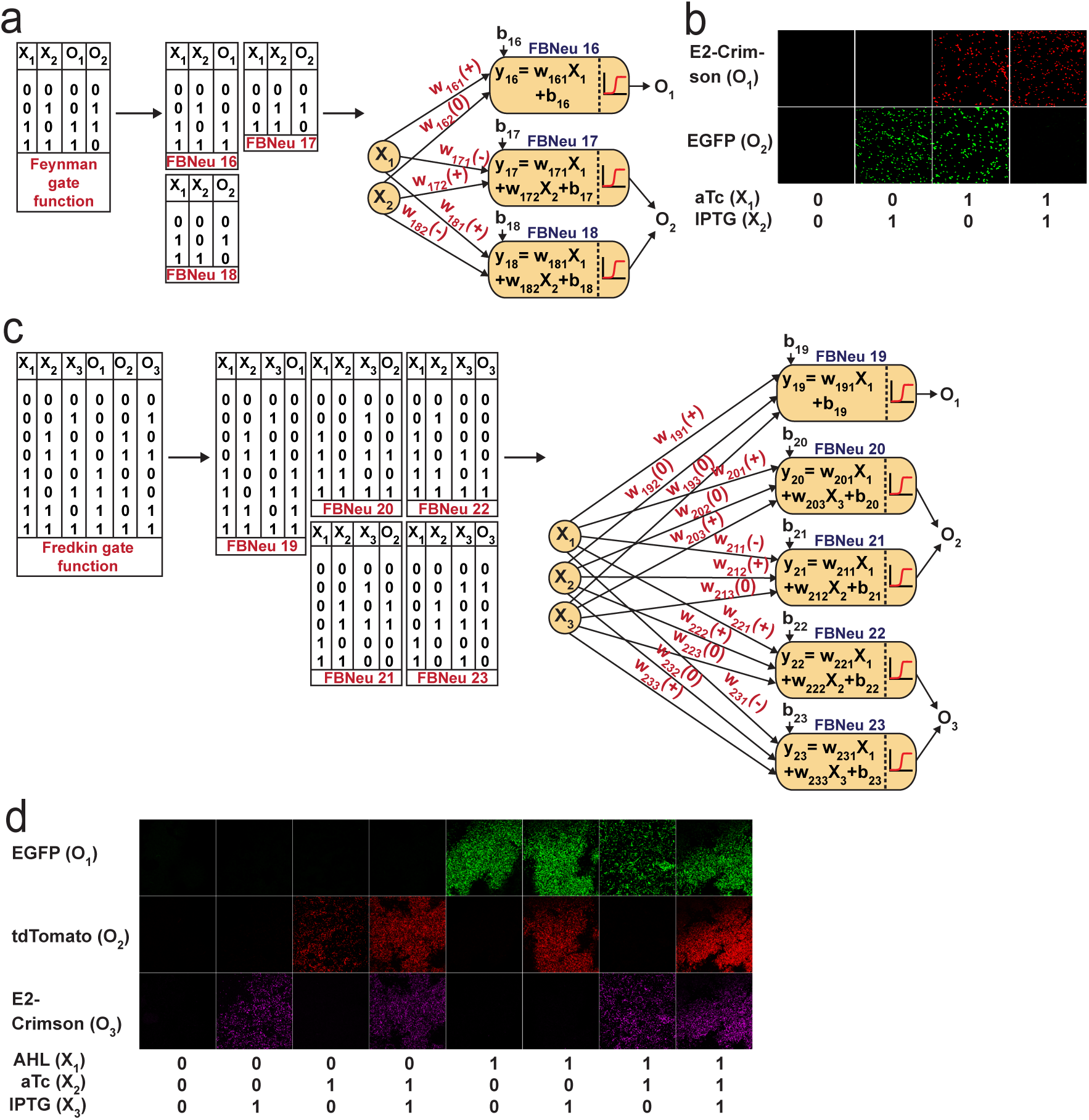
ANN Architechture and experimental demonstration of reversible Feynman and Fredkin gate with molecular engineered bactoneurons. Derivation of the functional bactoneurons from truth tables for **a)** Feynman gate and **c)** Fredkin gate. Experimental behavior of **b)** Feynman gate and **d)** Fredkin gate.

### Plausible correlations between ANN parameters and molecular mechanism

The abstract architectures and mathematical equations of an ANN are bridged with the physical bactoneurons through the molecular devices. Within the molecular devices, it may be possible to envisage the events of different inducers binding to the cognate transcription factors and causing conformational changes in them as a “summing unit” (linear combinations). The sensitivity of the transcription factors to their respective inducers can be seen in parallel to the synaptic strength of a neuron that is weights. The “summed” changes in the relevant transcription factors of a promoter directly affects their promoter binding capabilities and thereby modulates the activation of the promoter to express a gene. This can be perceived as the “activation unit”. In this work we manually adjusted the weights and bias of a neuron by changing the numbers and positions of various promoter operator sites, relative molecular amounts of transcription factors and promoter copy numbers through different plasmid copy numbers or altering RBS strength. We further showed a direct relation between ‘bias’ of an artificial neuron and leakage from the genetic devices. In this way, we established a clear correlation between ANN parameters and molecular mechanism of an intracellular molecular device. The weight adjustment was done manually by molecular engineering and the inference network was ‘uploaded’ in the final bactoneural networks. This might be compared with the systems, where the weight adjustments were done in the server and the inference network was sent to the mobile device^21^. It is important to note that the mathematical models for conventional bio-circuit design is different and highly specific for different functions and in fact, the mathematical model of a computing function may not guide the actual genetic circuit design^22^. Whereas, our ANN based frameworks required only single type of mathematical equation (Table S1) for creating different computing functions and we established a direct correlation between the sign of the weights in the activation function equation and nature of molecular mechanism (Fig. 1B). Thus, this allowed a simple and streamlined way to design molecular device for bactoneurons of any function directly guided by the mathematical equation of the activation function.

## Conclusions

In summary, we showed that the basic concept of ANN could be adapted in living bacteria with the help of molecular devices. The ANN type architecture with molecular engineered bacteria works as a flexible and general framework in its design and architecture for performing complex bio-computation, both conventional and reversible. Unlike, the conventional way of designing in-vitro small molecule computing^23^, enzyme based computing^24^, DNA computing^25^ and in-vivo synthetic genetic computing^26,27^, which followed the hierarchical logic circuit approach^28,29^, we showed that the ANN type framework can adapt a new design path to create complex computing function like encoder, majority function, mux, demux, Feynman gate and Fredkin gate, where encoder, majority function and reversible gates were demonstrated for the first time in living cells. Reversible computing is a new class of computing for biological cells and our work might pave the way in that direction. In ANN, any function can be designed and simulated just by adjusting the weights and bias values of a single mathematical equation and we established a direct relation between signs of the weights and nature of the interaction in the molecular devices. Thus, our approach established a new streamlined design and construction platform complementary to the conventional bio-circuit design^22^ and may have implications in complex reversible and irreversible biocomputing, bacteria-based ANN hardware, and synthetic biology.

## Materials and Methods

### Promoters and genes, plasmids, RBS’s & primers

The genetic devices were made according to the designs. The bioparts (promoter, ribosome binding sites (RBS), gene and transcription terminators) were arranged within appropriate plasmids using standard molecular biology protocols. The designed promoters were obtained by either PCR (KOD Hot-Start DNA polymerase, Merck-Millipore). PCR amplification was performed by KOD Hot Start DNA polymerase (Merck Millipore). All enzymes, ligase, and ladders were from New England BioLabs. Plasmid isolation, gel extraction, and PCR purification kits were from QIAGEN. Translation initiation rate for EGFP and CI under the control of promoter P_AAH_ and P_LtetO-1_/P_LlacO-1_ respectively was calculated and weak RBSs (RBSs RH and RC1-3) were designed using RBS Calculator v2.0^42^, considering RBS R (BBa_B0034)^43^ along with linker GGTACC (*Kpn*I site) as degenerate RBS sequence and E. coli-MG1655 as the organism. All promoter sequences, RBS sequences, primers, and plasmids are shown in supplementary tables S3, S6, S8, and S9 respectively. All primers, oligos and gene products were synthesized from IDT and Invitrogen. All cloned genes, promoters, RBS’s in plasmid constructs were sequence verified by Eurofins Genomics India Pvt. Ltd., Bangalore, India.

### Bacterial cell culture for characterization

Chemically competent *Escherichia coli* DH5α strain was used for cloning and DH5αZ1 strain was used for the experimental characterization. Working concentrations of the antibiotics in LB-Agar, Miller (Difco, Beckton Dickinson) plates as well as in LB broth, Miller (Difco, Beckton Dickinson) were: 100 μg/ml for Ampicillin (Himedia), 34 μg/ml for Chloramphenicol (Himedia) and 50 μg/ml for Kanamycin (Sigma Aldrich). DH5αZ1 cells were transformed with appropriate Sequence verified plasmid constructs. Well-isolated single colonies were picked from LB agar-plates, inoculated to fresh LB-liquid media, and grown overnight in presence of antibiotics. Next, the overnight culture was re-diluted 100 times in fresh LB media with antibiotics and with or without inducers as per the design of the gene circuit, and grown at 37° C, ~ 250 rpm. Engineered cells for weight and bias adjustment steps (expression characterization and dose response experiments) were grown for 16 hours and cells with final constructs were grown for 10 hours with inducers, resuspended and grown for another 6 hours. These 10+6 hours growth was performed for expression characterization, dose response experiments, validation experiments and for full ANN microscopy experiments.

### Dose-response experiments and validation experiments

All dose-response experiments were performed by varying one inducer across 9 or more concentration points while the other inducer was kept constant (“0” state or “1” as the case may be). Here, for the linear combinations of input signals, we converted the concentration range of each input chemical from ‘0’ to ‘1’, where 0 signifies zero concentration and 1 signifies the saturating concentration of the chemical. Any concentration higher than the saturation concentration was treated as 1. For the validation experiments, the corresponding two inducers of the relevant constructs were simultaneously varied across 10+ concentration points. For, validation experiments, we used different concentration points compare to the concentration used in dose response. For intermediate experiments during weight and bias adjustments of the constructs, single colony was used. For the final constructs, all experiments data from minimum 3 independent colonies were collected.

### Measurement of fluorescence and optical density, Normalization, and Scaling

For fluorescence and optical density (OD) measurements, Synergy HTX Multi-Mode reader (Biotek Instruments, USA) was used. For this purpose, cells were diluted in PBS (pH 7.4) to reach around OD600 as 0.8, loaded onto 96-well multi-well plate (black, Greiner Bio-One), and both EGFP fluorescence with appropriate gain and OD600 was measured. For EGFP fluorescence measurements, we used 485/20 nm excitation filter and 516/20 nm emission bandpass filter. At least 3 biological replicates had been considered for each condition to collect the fluorescence and OD data. The raw fluorescence values were divided by respective OD600 values and thus normalized to the number of cells. Auto-fluorescence was measured as average normalized fluorescence of the untransformed DH5αZ1 set (no plasmid set) and subtracted from the normalized fluorescence value of the experimental set. The above normalization can be mathematically represented as follows: –

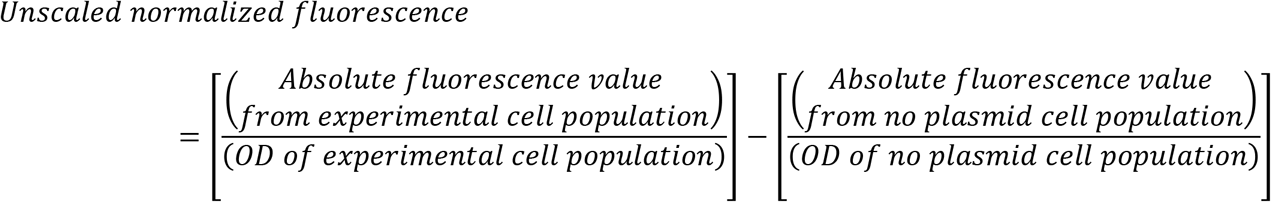

The values thus obtained were then scaled down between 0 and 1-considering the normalized fluorescence value at the induction point of maximum expected fluorescence to be 1,

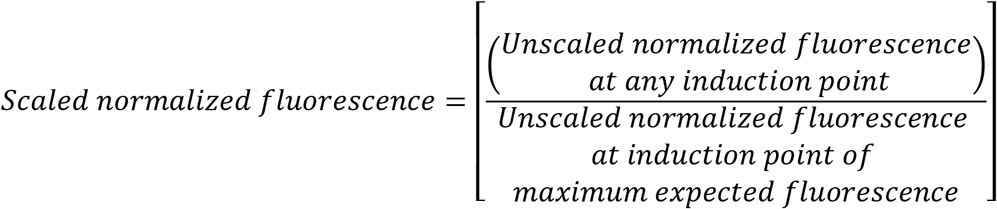

### Data Analysis, Fitting, Mathematical Modelling and Simulation

The fittings of the dose response curves were done by appropriate equations (Supplementary Table S10).

However, all the equations could be brought down in a general form,

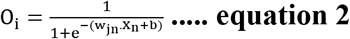

where,

O_i_ is the output signal from the artificial neuron j,

X_n_ corresponds to the magnitude of the varying input (n) to the artificial neuron j,

w_n_ corresponds to the weight of the n^th^ input to the artificial neuron j, and, b is given by:

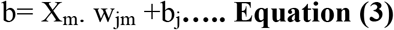

where,

X_m_ corresponds to the logical state of the constant input (m) to the artificial neuron j (0 or 1),

w_m_ corresponds to the weight of the m^th^ input to the artificial neuron j,

b_j_ corresponds to the bias of the artificial neuron j.

The scaled output fluorescence values obtained from the dose-response experiments were plotted against the varying inducer concentration and fitted against equation 2. All data analysis and fitting were performed in OriginPro 2018 (OriginLab Corporation, USA) and was performed using built-in Levenberg Marquardt algorithm, a damped least squares (DLS) method. The parameter “w_jn_” of the fitting function (equation 2) gives the “weight” of the varying input in the summation function of the corresponding neuron j. The b value obtained includes the bias plus the product of the input logic state of the second input and its weight as explained above. Upon similarly fitting the dose-response of the neuron to the second input, the weight “w_j_” and “b” for input 2 is obtained. Solving equation 2 and 3 for both the inputs, “w_jinput1_”, “w_jinput2_” and “b_j_” of the complete summation function of the corresponding neuron j was obtained. All parameter values for all constructs were shown in supplementary table S4. The simulations were performed by generating matrices of calculated normalized output fluorescence values against simultaneously varying concentrations of the corresponding two inputs across 65X65 or more points following the parameterized activation function. For single-input systems, simulation for a given activation function was carried out across 19 varying input concentration points.

### Microscopy

DH5αZ1 cells were transformed with the appropriate sequence-verified plasmid construct(s). Following an 10 hours induction followed by re-suspension and 6 hour induction step, cells were washed thrice in PBS. Cell pellets were finally resuspended in fresh PBS (pH 7.2-7.4) and this re-suspension was used to prepare fresh slides. A Laser Scanning Microscope Zeiss LSM 710/ ConfoCor 3 operating on ZEN 2008 software was used for imaging of the de-multiplexer, multiplexer, majority function, decoder and encoder. The cell suspension slides were subjected to excitation by appropriate laser channels (458 nm Ar Laser for mTFP1, 488 nm Ar Laser for EGFP, 514 nm Ar Laser for mVenus, 543 nm He-Ne Laser for mKO2 and 633 nm He-Ne Laser for E2-Crimson) and fluorescence emissions were captured through proper emission filters (BP484-504 nm for mTFP1, BP500-520 nm for EGFP, BP521-541 nm for mVenus, BP 561-591 nm for mKO2, BP641-670nm (2-to-4 decoder)/BP630-650 nm (1-to-2 de-multiplexer) for E2-Crimson) with a 63x Oil immersion objective and were detected through a T-PMT. The pin hole was completely open. For reversible Fredkin gate imaging, Nikon AIR si confocal microsocpe along with resonant scanner and coherent CUBE diode laser system was used. The mixed cell populations, washed and resuspended in PBS (pH 7.2-7.4), was added on the top of the 1% molten agarose pad which was placed upon a cleaned glass slide. The sample field was then covered with clear cover slip, placed under 60 X water immersion and subjected to excitation by laser channels (488 nm Laser for EGFP, 561 nm for td-Tomato and 640 nm for E2-Crimson). Three different emission filters (BP 525/50 nm for EGFP, BP 585/65 for td-Tomato and BP 700/75 nm for E2-Crimson) were used for collecting fluorescence and differential interference constrast (DIC) images were captured for all samples as well. Microscopic images were processed through ImageJ software for better visualization.

## Acknowledgements

This work was financially supported by SINP intramural funding (Department of Atomic Energy, Govt. of India), SERB (CRG/20lB/00l394), and Ramanujan Fellowship (DST), Govt. of India. We thank Mr. Sayak Mukhopadhyay for constructing two plasmids pA2MCS and pC3MCS.

## Author Contributions

SB conceived and designed the study. KS, DB, and RS performed all the experiments. KS, DB, and SB designed the experiments, analyzed and interpreted the data, and wrote the paper.

## Competing Interests statement

We have no competing interest.

## Supplementary Information

**Figure S1:**
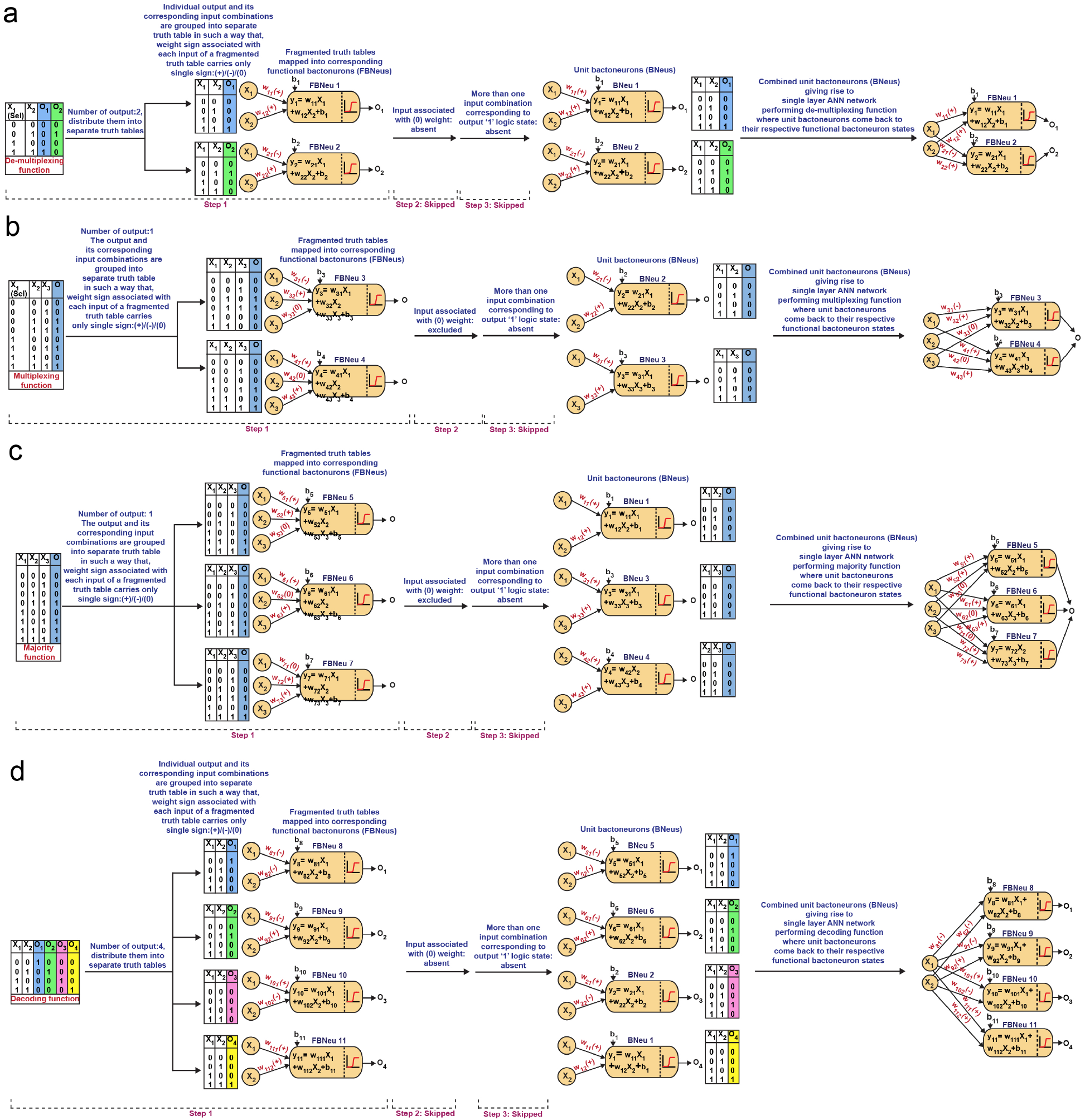
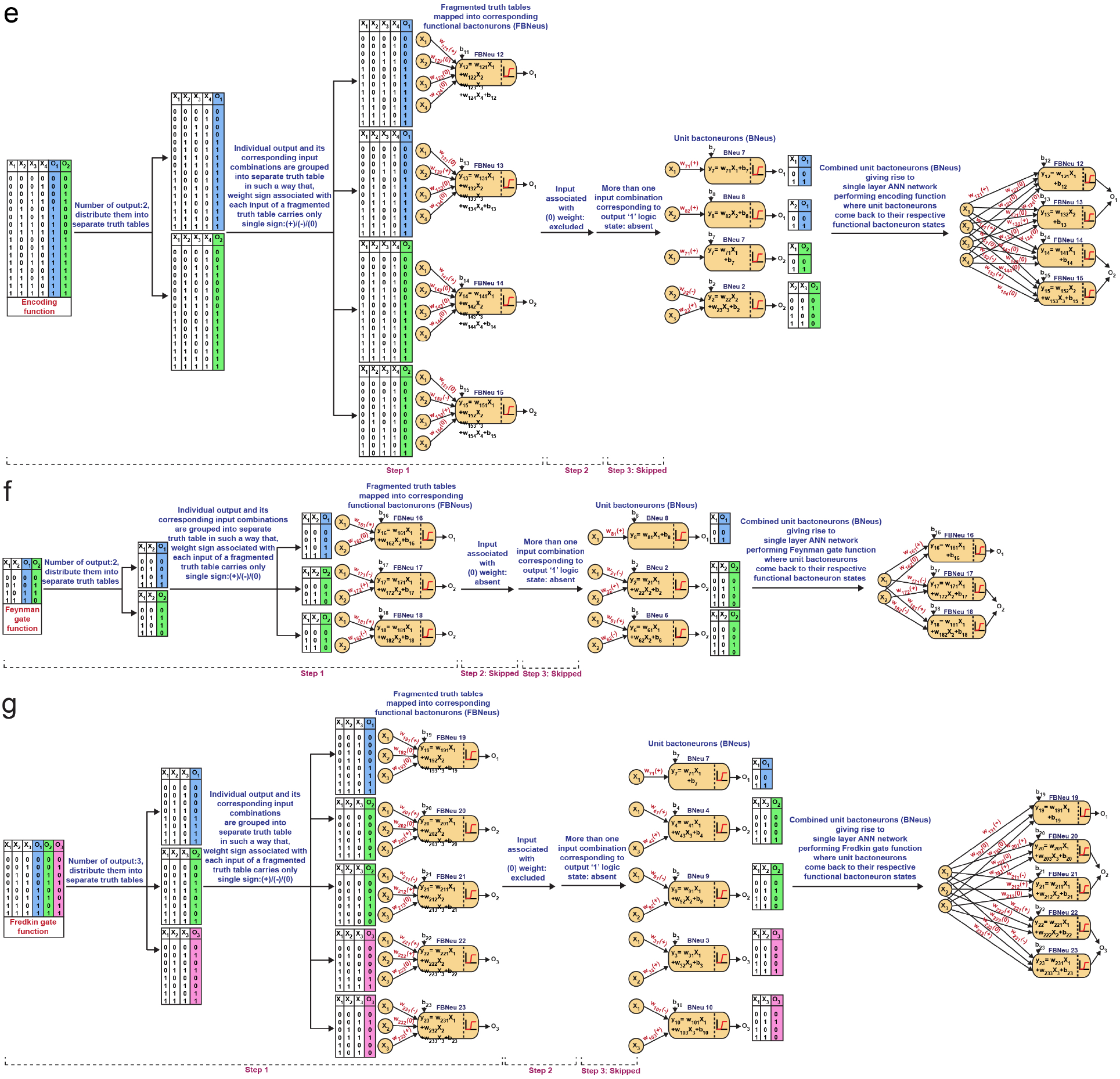
Derivation of functional and unit bactoneurons for a) de-multiplexing function, b) multiplexing function, c) majority function, d) decoding function, e) encoding function, f) Feynman gate function and g) Fredkin gate function. In each case, combination of unit bactoneurons gives rise to single layer ANN architecture where, individual unit bactoneurons come back to their corresponding functional bactoneuron states while they get associated with ‘0’ weighted inputs (If any). In the network level, parts of the summation function of each functional bactoneuron, contributed by ‘0’ weighted input, are not shown.

**Figure S2:**
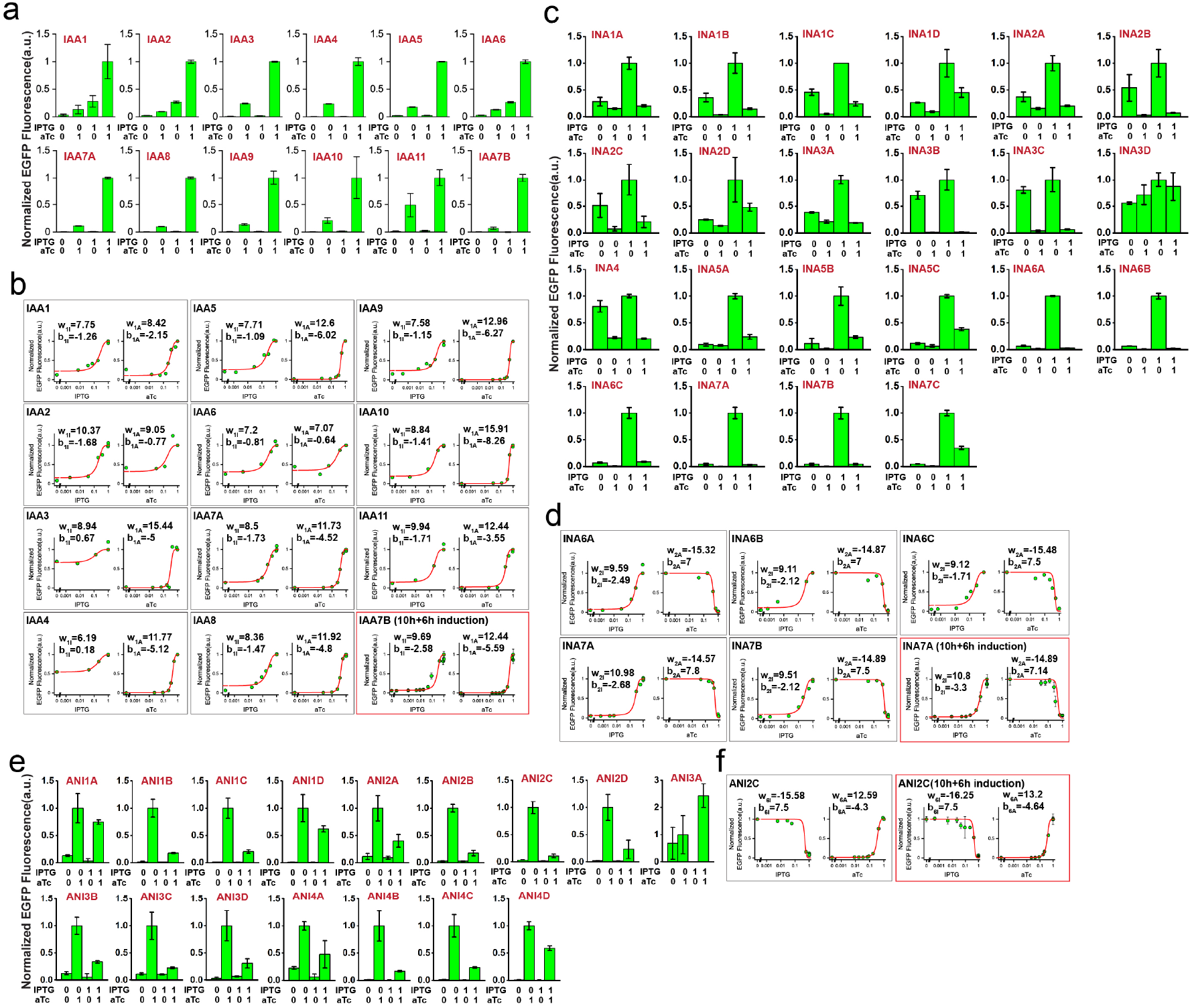
Details of characterization and dose responses of BNeus 1, 2 and 6. Expression characterization of **a)** BNeu 1, **c)** BNeu 2 and **e)** BNeu 6 and dose responses of **b)** BNeu 1, **d)** BNeu 2 and **f)** BNeu 6.

**Figure S3:**
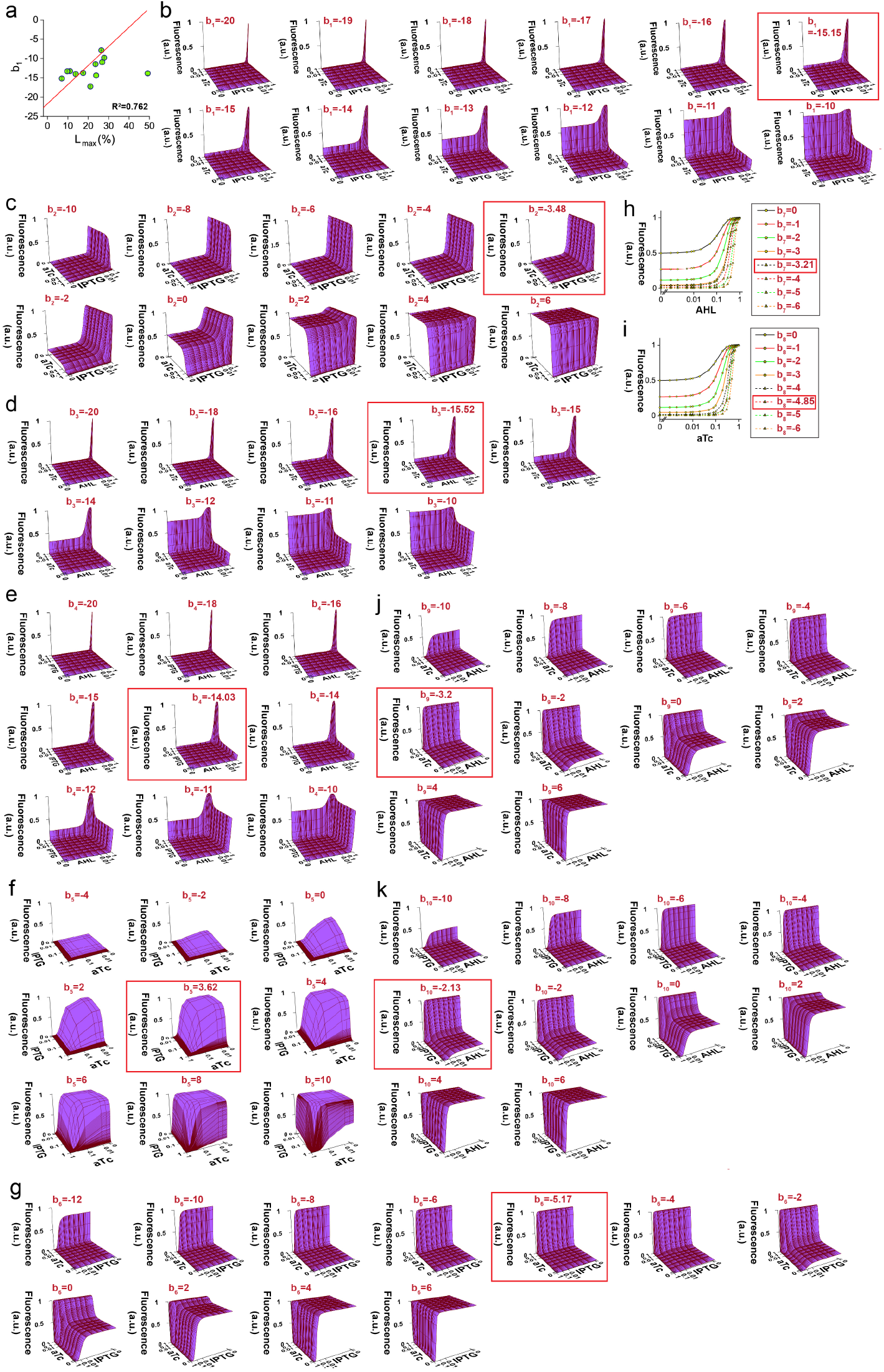
Correlation between bias and leakage of a Unit bactoneuron. **a)** Correlation between bias (b_1_) and the percentage highest leakage (L_max_ (%)) for all BNeu 1 cellular devices obtained from weight and bias adjustment steps. Simulated output behaviors of **b)** BNeu 1, **c)** BNeu 2 **d)** BNeu 3 **e)** BNeu 4, **f)** BNeu 5, **g)** BNeu 6, **h)** BNeu 7, **i)** BNeu 8, **j)** BNeu 9 and **k)** BNeu 10. Simulation corresponding to the bias value obtained experimentally for each bactoneuron is shown in red box.

**Figure S4:**
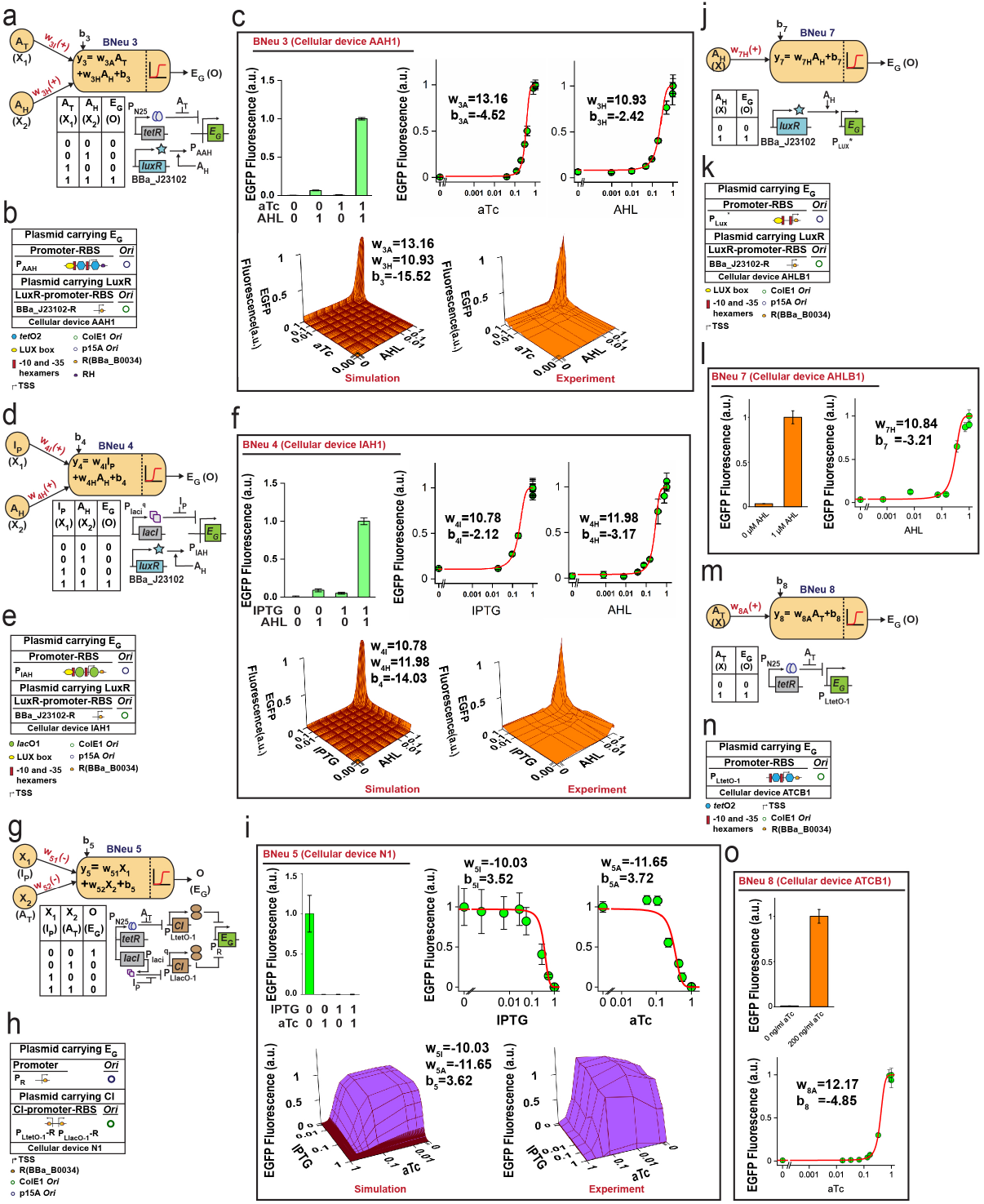
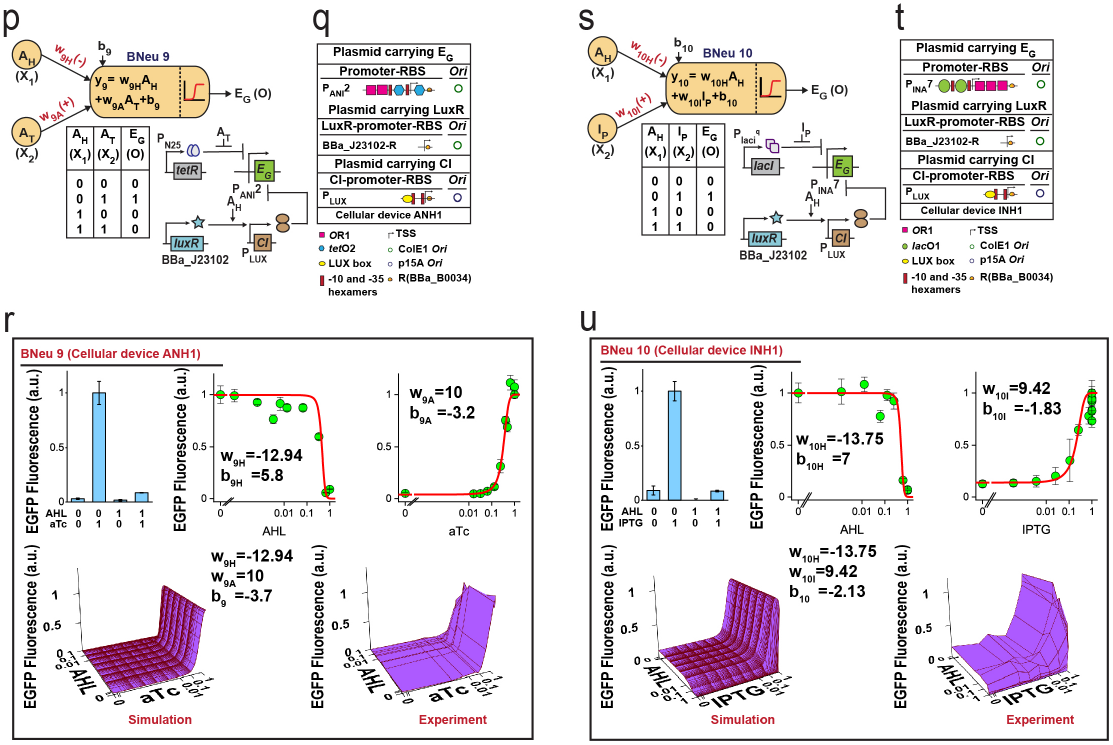
Characterization of unit bactoneurons BNeu 3, BNeu 4, BNeu 5, BNeu 7, BNeu 8, BNeu 9 and BNeu 10. Neural architectures, truth tables and biological circuit designs of unit bactoneurons **a)** BNeu 3, **d)** BNeu 4, **g)** BNeu 5, **j)** BNeu 7, **m)** BNeu 8, **p)** BNeu 9 and **s)** BNeu 10 are shown. Details of plasmids carrying bioparts of the biological circuit designs of **b)** BNeu 3, **e)** BNeu 4, **h)** BNeu 5, **k)** BNeu 7, **n)** BNeu 8, **q)** BNeu 9 and **t)** BNeu 10 are illustrated. Expression characterization, dose responses, 3D simulations and experimental 3D behavior of **c)** BNeu 3, **f)** BNeu 4, **i)** BNeu 5, **l)** BNeu 7, **o)** BNeu 8, **r)** BNeu 9 and **u)** BNeu 10 in terms of EGFP expression are also shown.

**Figure S5:**
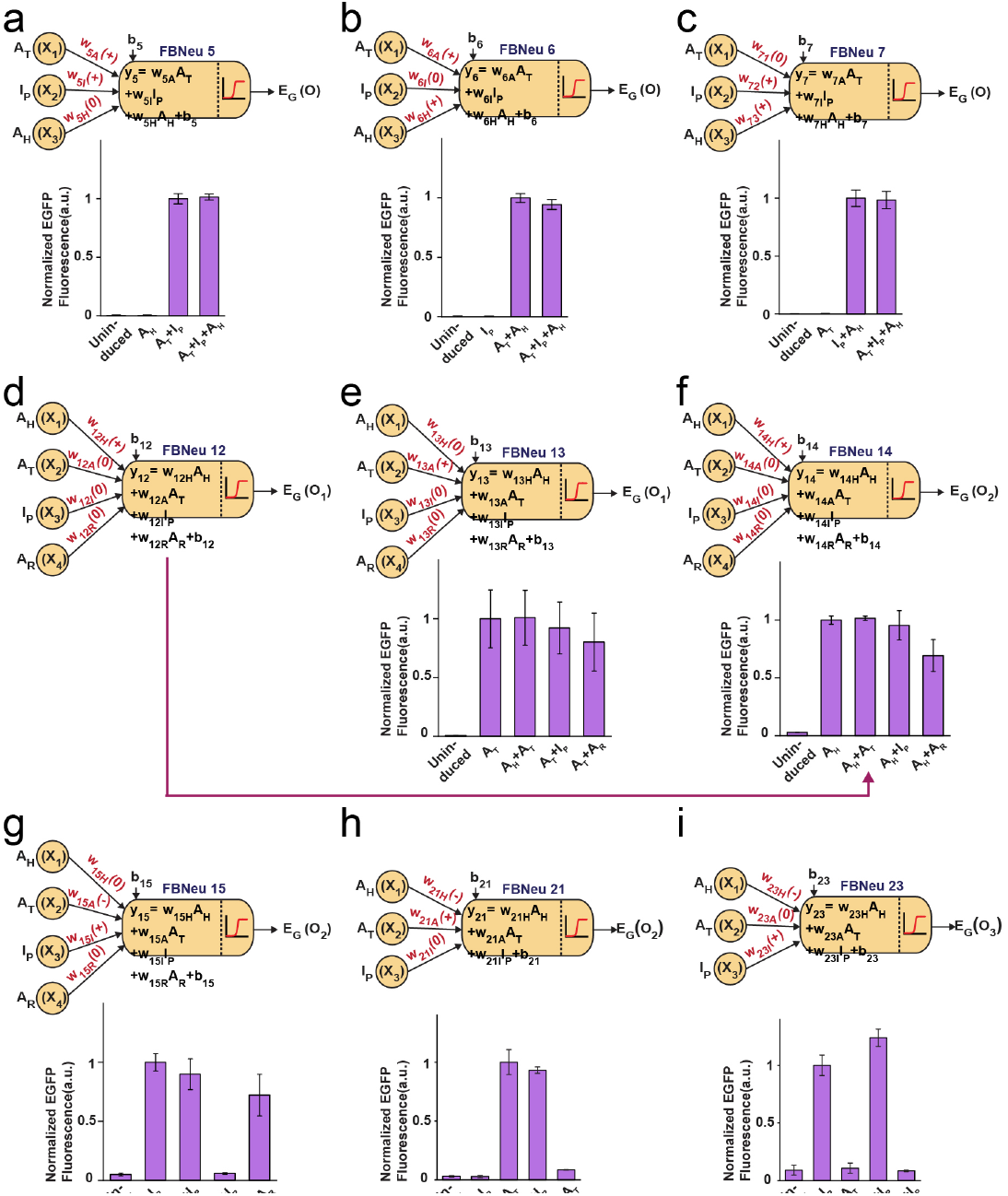
Characterizations of functional bactoneurons associated with weight=0 towards specific inducers. Each functional bactoneuron population was subjected to 10h+6h induction with a set of inducers which was chosen based on the neural architecture of individual functional bactoneurons, and then characterized in terms of EGFP expression. If the presence or absence of a specific inducer didn’t change the output of the functional bactoneuron, then only we considered that, the inducer was associated with zero weight. Neural architectures and validation for ‘0’ weighted input(s) of functional bactoneurons **a)** FBNeu 5, **b)** FBNeu 6, **c)** FBNeu 7, **d)** FBNeu 12, **e)** FBNeu 13, **f)** FBNeu 14, **g)** FBNeu 15, **h)** FBNeu 21 and **i)** FBNeu 23 are shown. FBNeu 20 and FBNeu 22 from Fredkin gate function are equivalent to FBNeu 7 and FBNeu 6 respectively except their different outputs. Therefore, individual weight ‘0’ input validation for FBNeus 6 and 7 justifies the same for FBNeus 22 and 20 as well. FBNeu 12 and FBNeu 14 are similar except their outputs. Here, individual functional bactoneurons are characterized in terms of EGFP output. Therefore, both FBNeu 12 and FBNeu 14 produce EGFP output and hence, they become identical. Thus, they share common ‘0’ weighted input validation data (Shown with magenta arrow). FBNeu 16 from Feynman gate function is a sub-set of FBNeu 13 as it operates on lesser number of inputs whereas, their corresponding unit bactoneuron is common. Therefore, weight ‘0’ input of FBNeu 16 can be validated from the characterization result of FBNeu 13. Similarly, weight ‘0’ inputs of Fredkin gate functional bactoneuron FBNeu 19 can be validated by characterization result of FBNeu 12/14.

**Figure S6:**
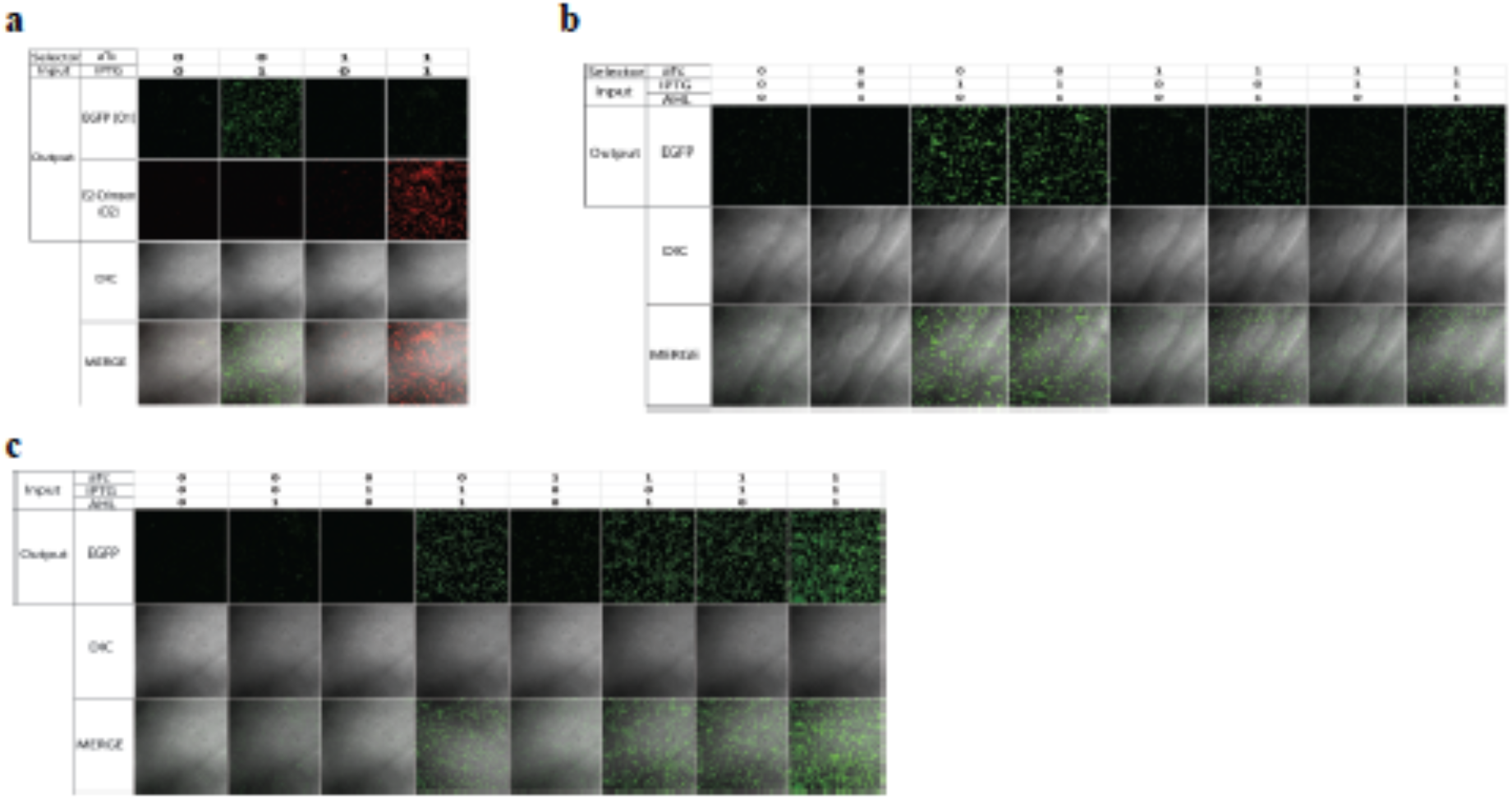
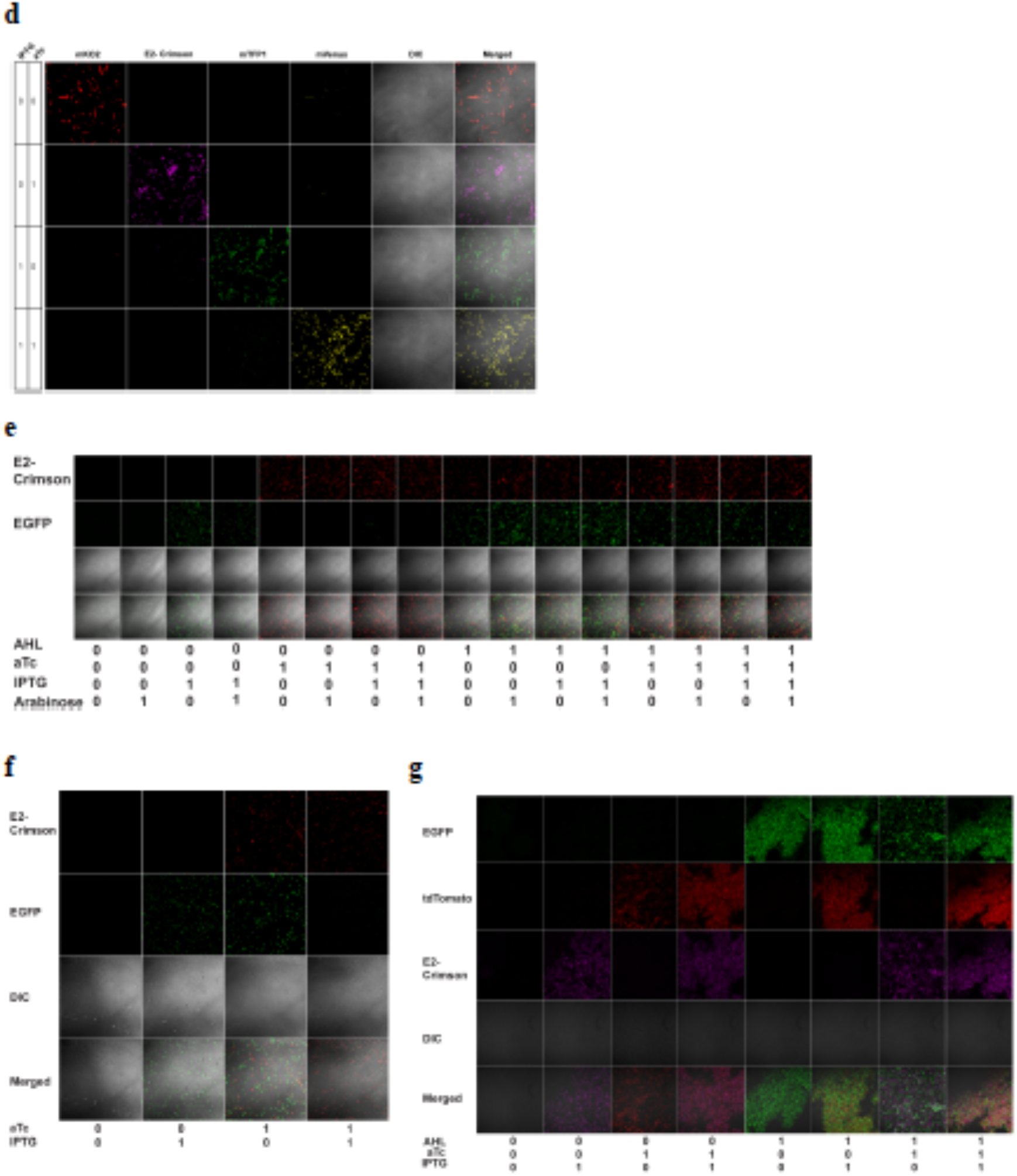
Microscopy images with corresponding differential interference contrast (DIC) and merged channels. Populations of different combinations of bactoneurons, depending on the complex functions they constitute, were co-cultured with appropriate inductions where, they together formed a bactoneural layer. They were viewed under relevant laser channels and emission filters. DIC images show a heterogenous population of cells in the field with each sub-population responding uniquely to the induction conditions. A bactoneuron’s activation is reported by fluorescence from its respective output protein whereas inactive bactoneurons show no fluorescence. Microscopic images for the bactoneuron-based single layer ANN type architectures for **a)** de-multiplexing function, **b)** multiplexing function, **c)** majority function, **d)** decoding function, encoding function **f**) reversible Feynman gate and **g)** reversible Fredkin gate are shown.

**Table S1:**
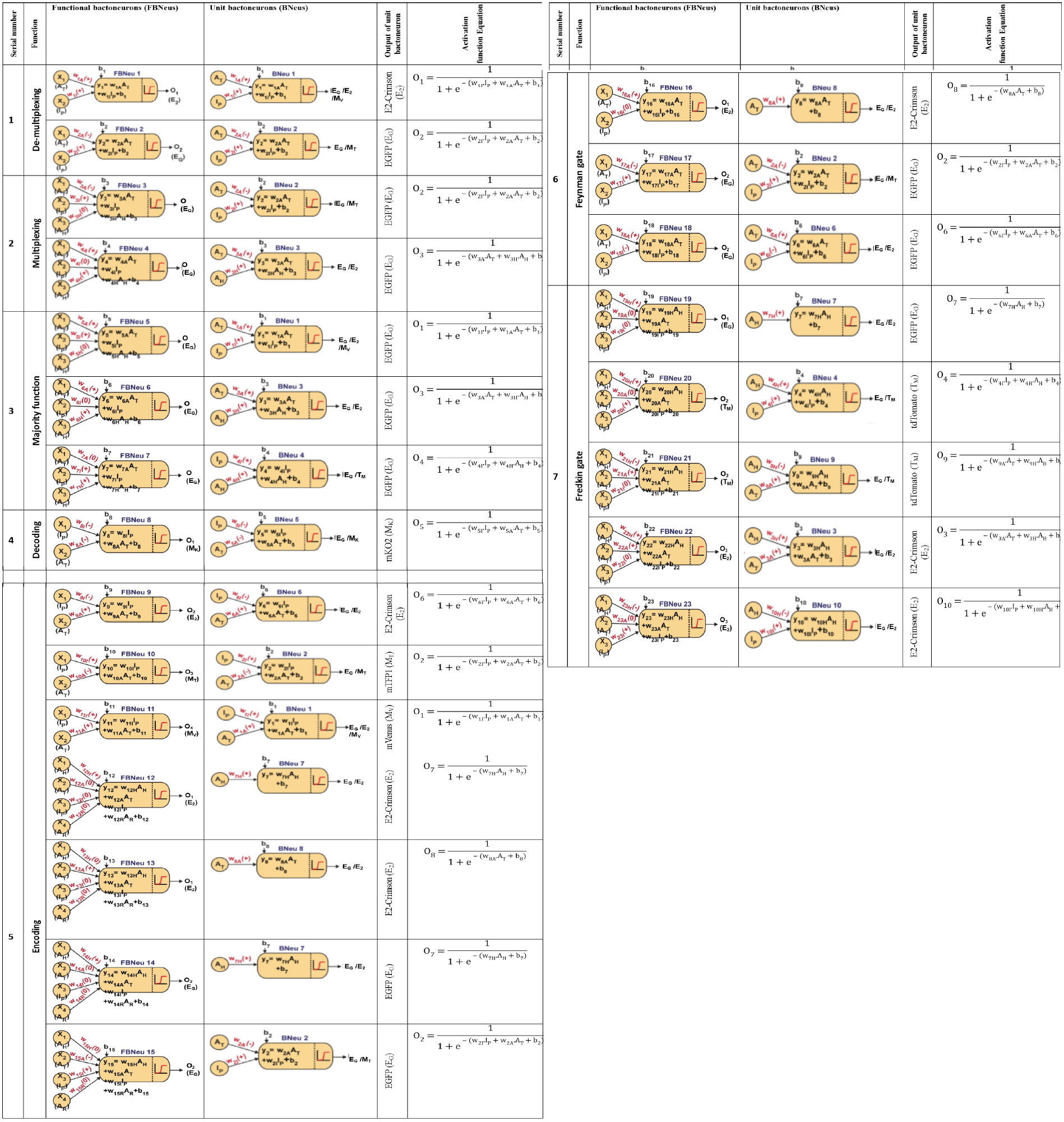
Details of functional bactoneurons and corresponding unit bactoneurons associated with the computing functions developed in this study. Output fluorescent proteins and activation function equations corresponding to unit bactoneurons are also described.

**Table S2:**
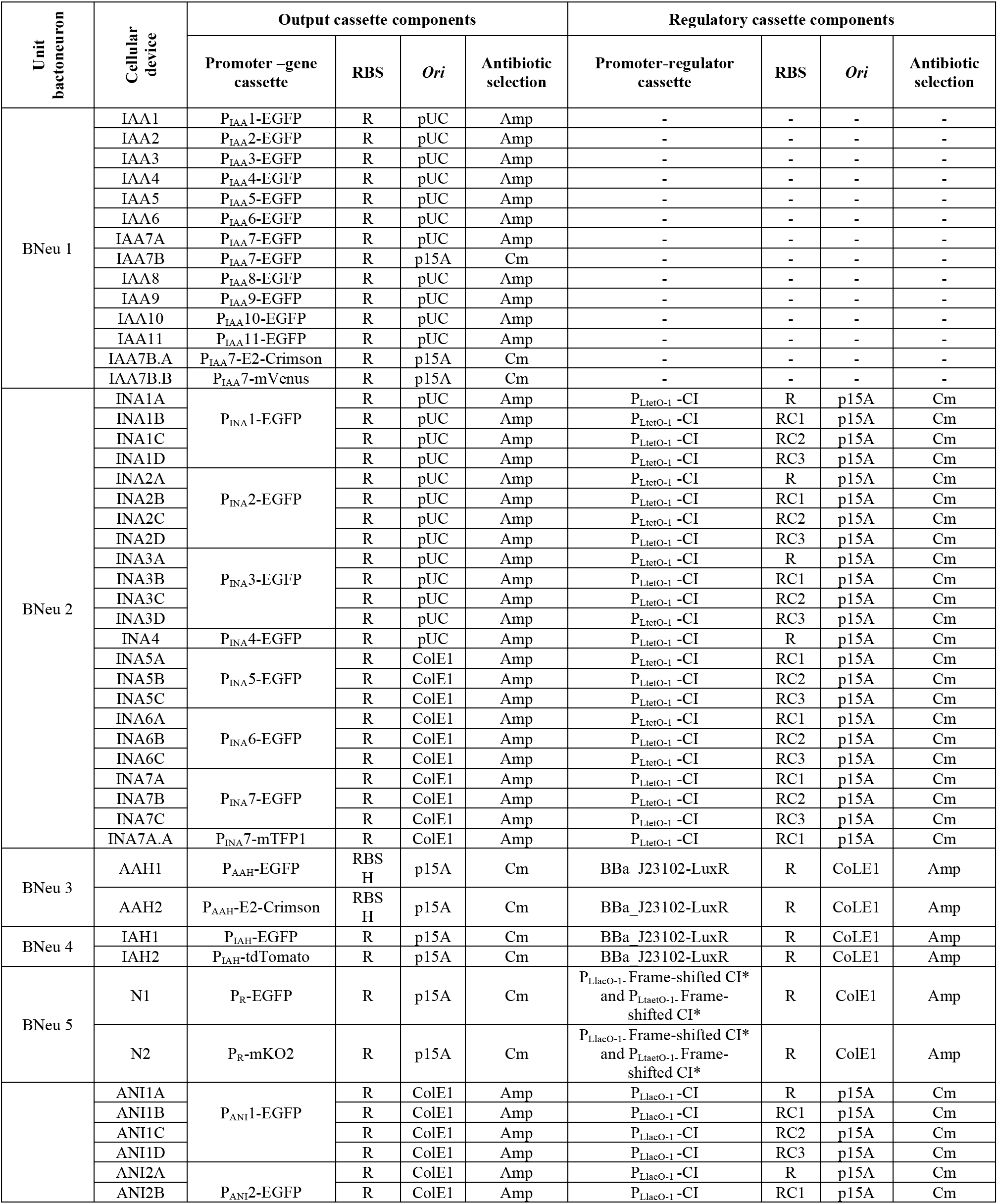
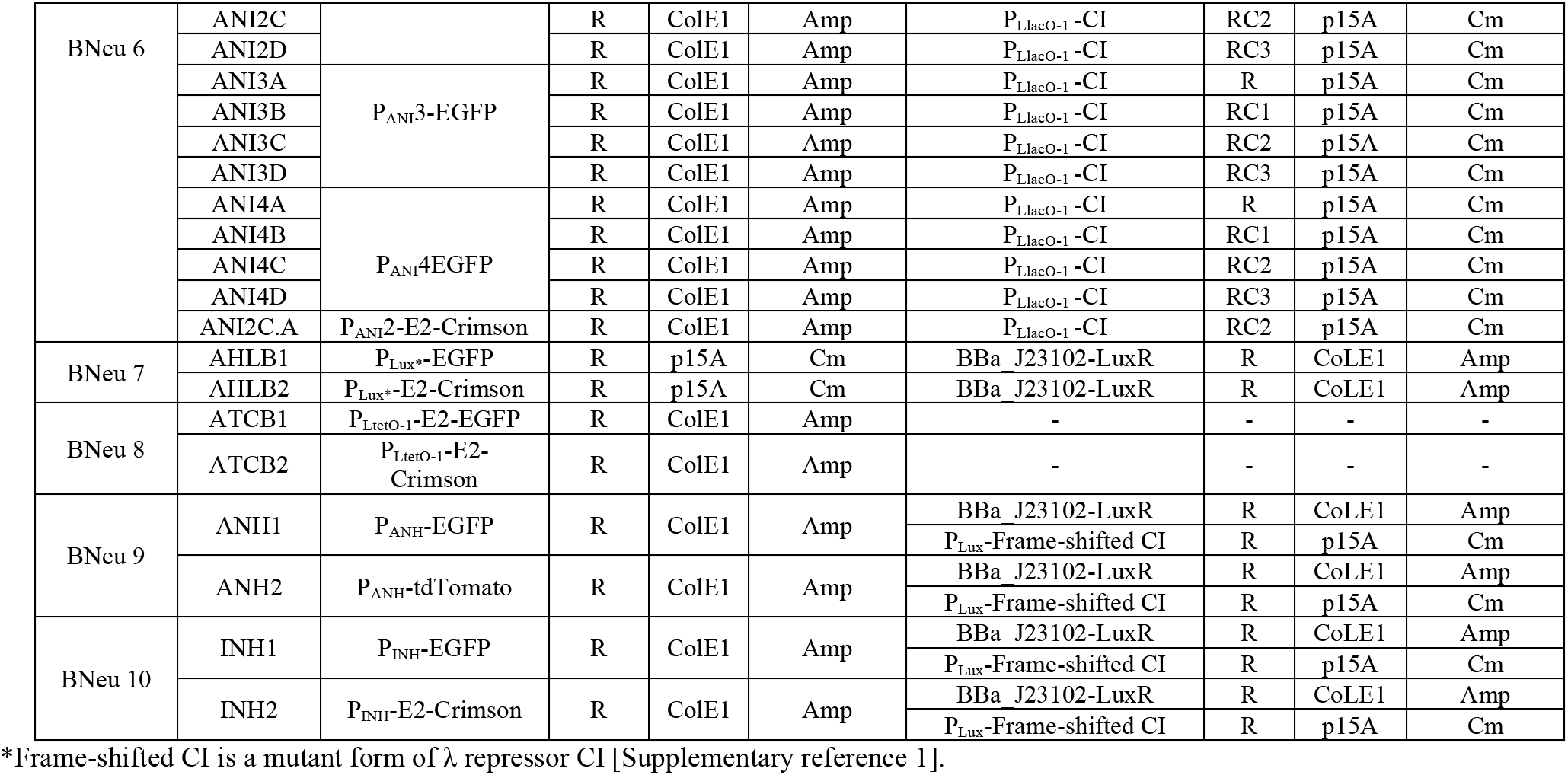
List of cellular devices constructed in this study.

**Table S3:**
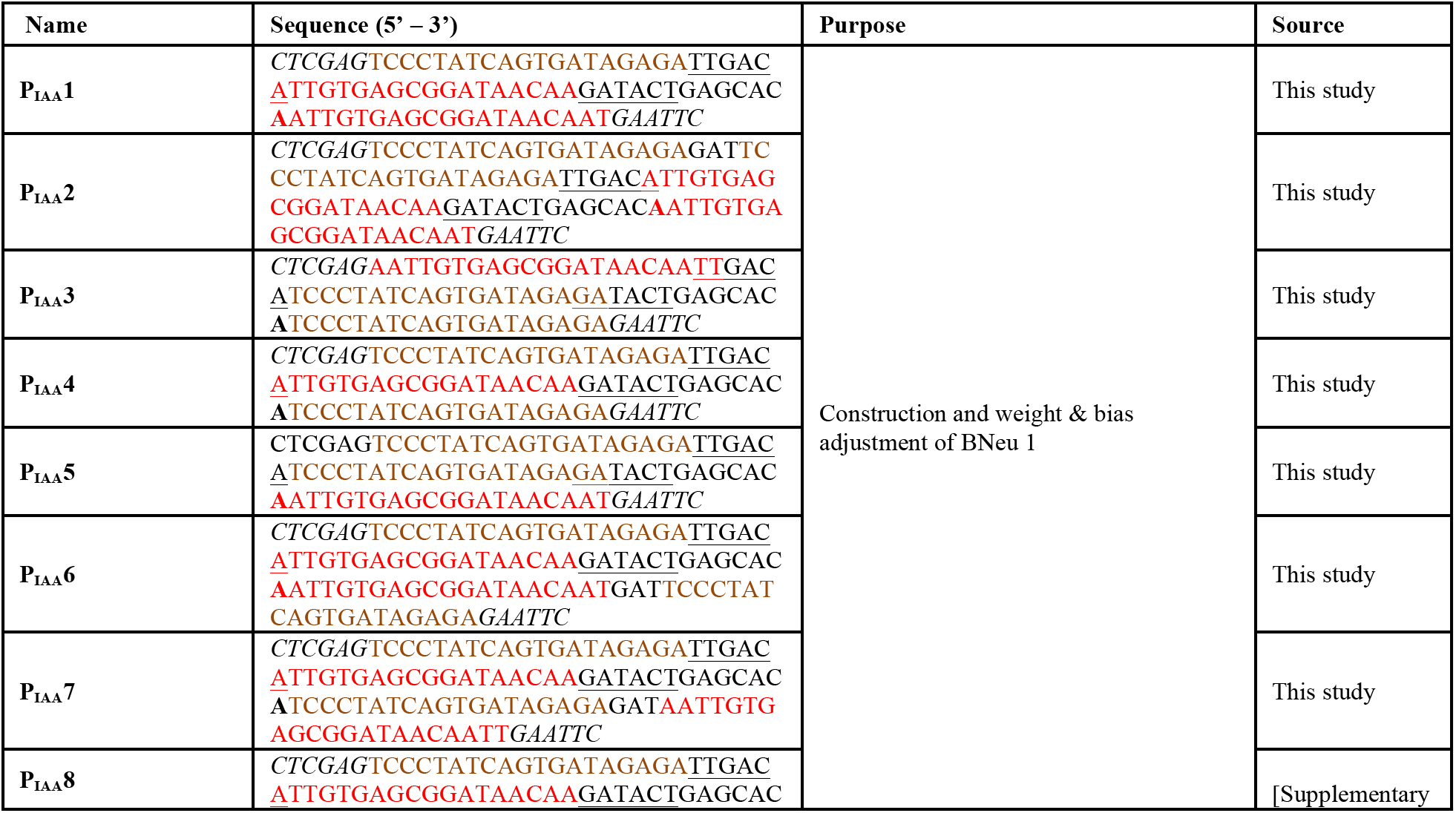
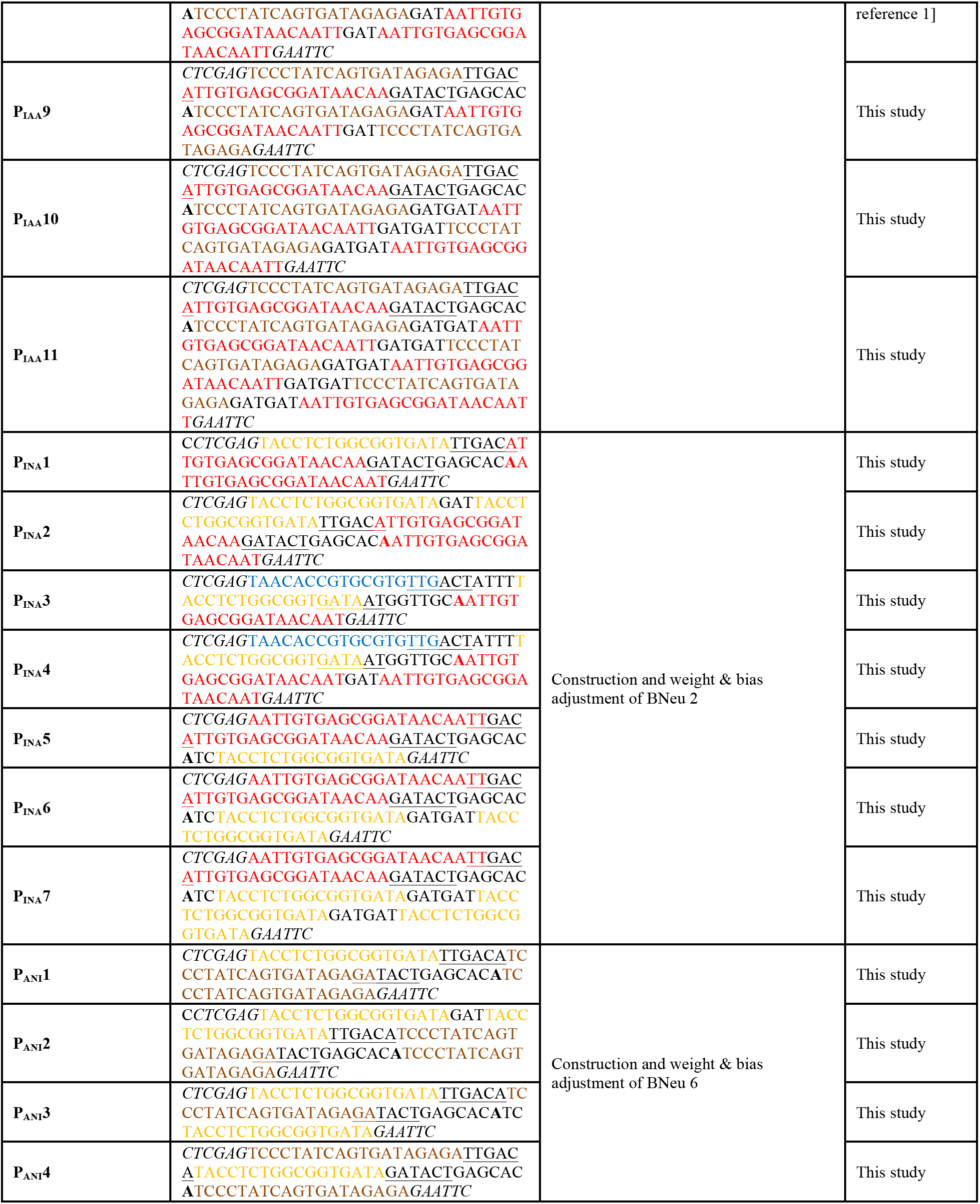
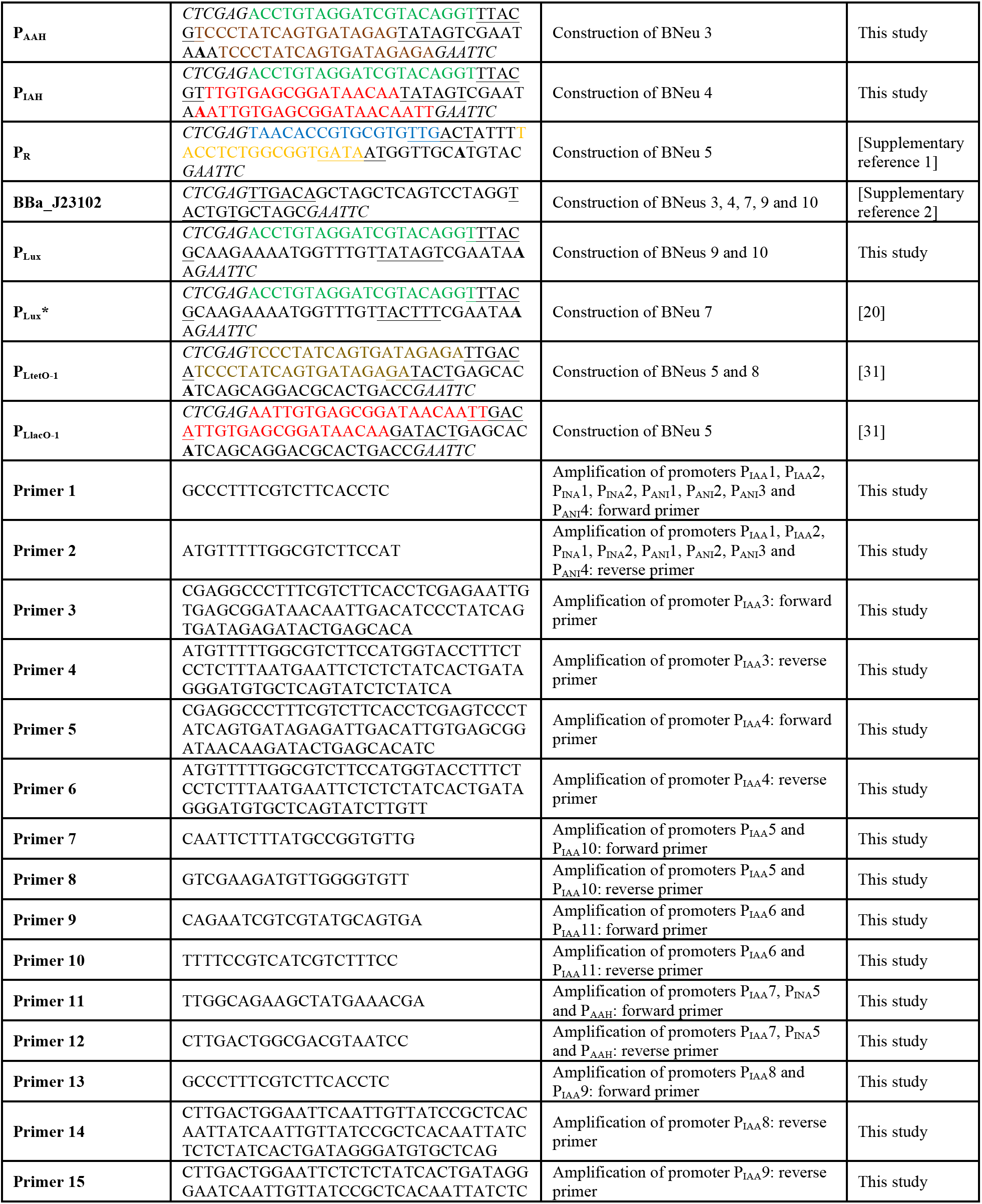
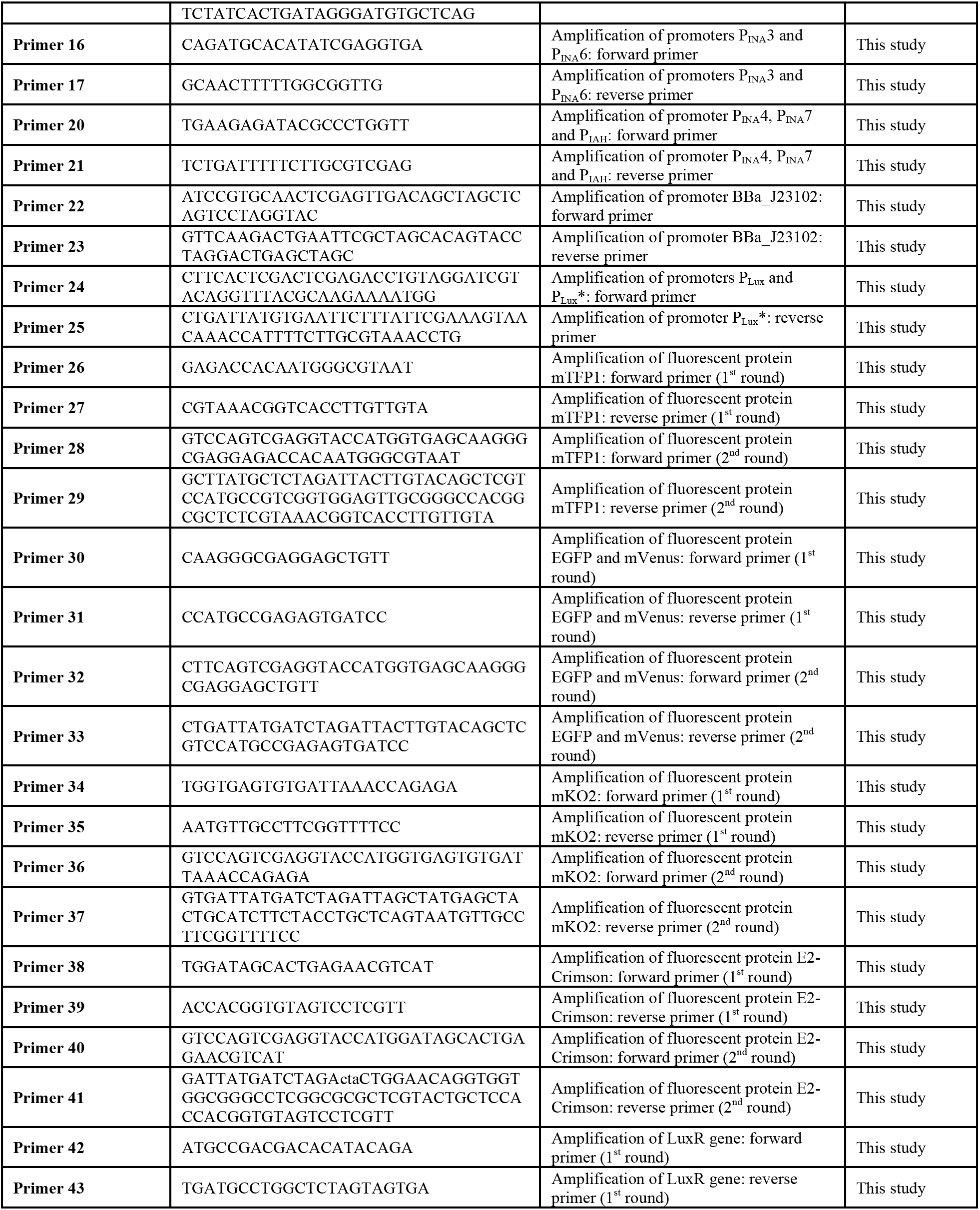
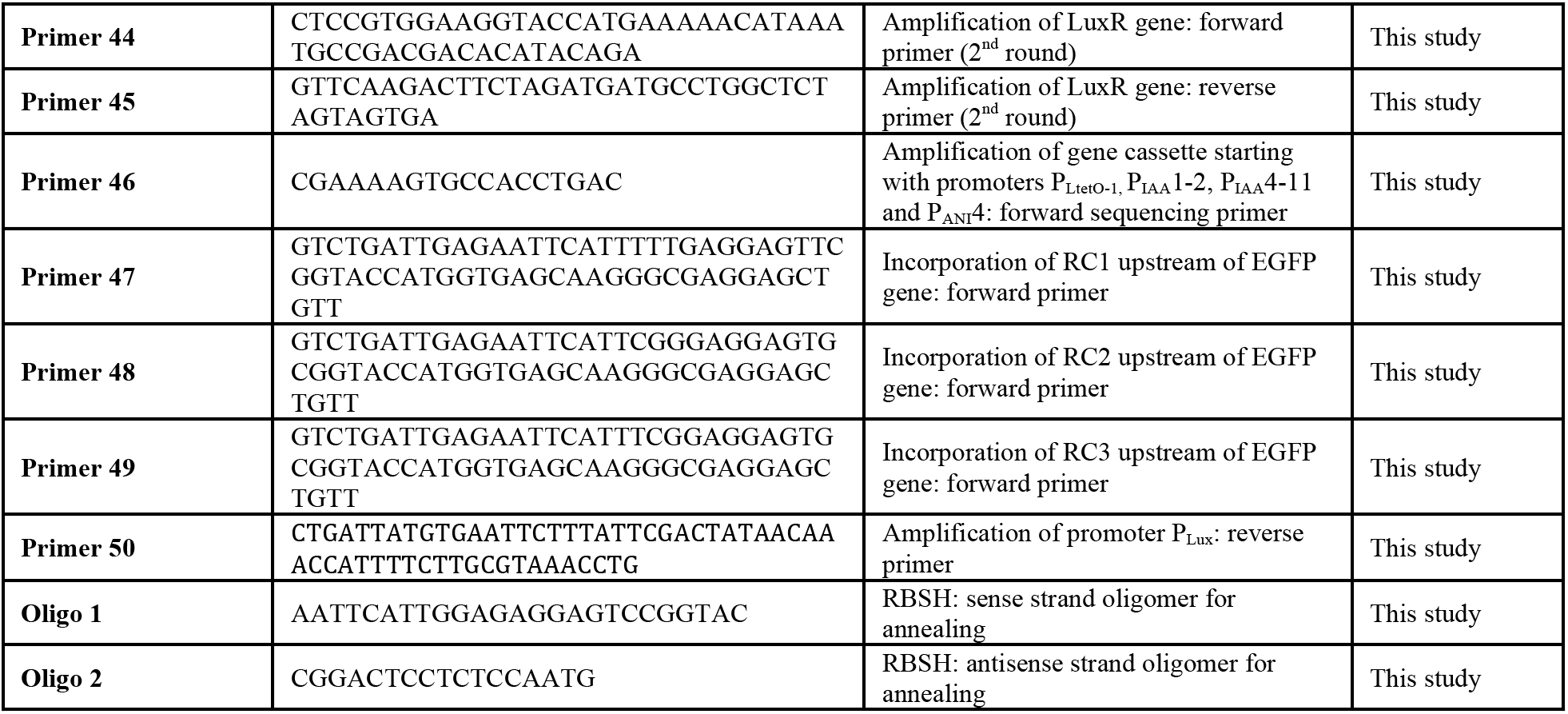
List of Promoters, primers, oligos and RBSs. *lac*O1, *tet*O2, Lux box, *O*R1 and *O*R2 operator sites are colored in red, brown, green, yellow and blue respectively. Transcription start site is shown in bold. −10 and −35 hexamers are underlined. Each promoter is flanked by *Xho*I and *EcoR*I restriction sites (marked in italics).

**Table S4:**
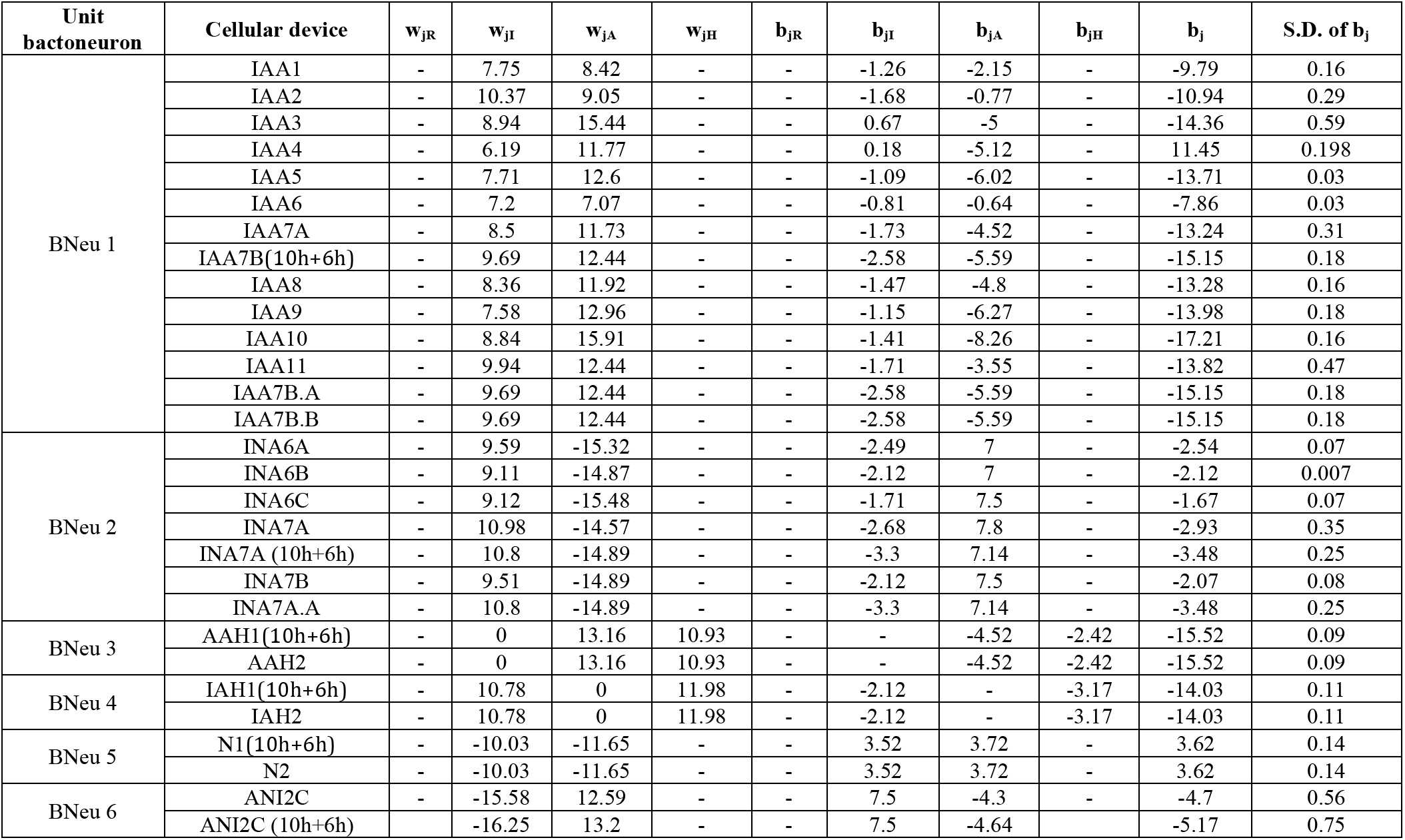
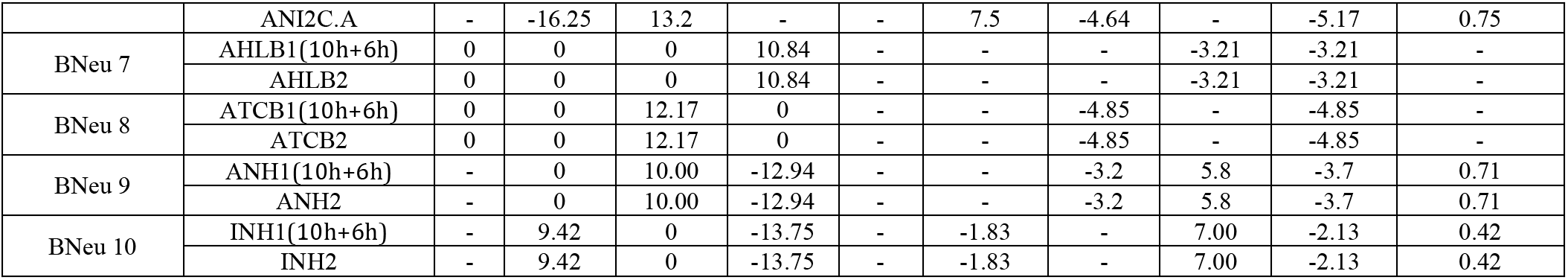
Weights and biases of each cellular device (construct) used for optimizing and improving corresponding unit bactoneuron (BNeu j).

**Table S5:**
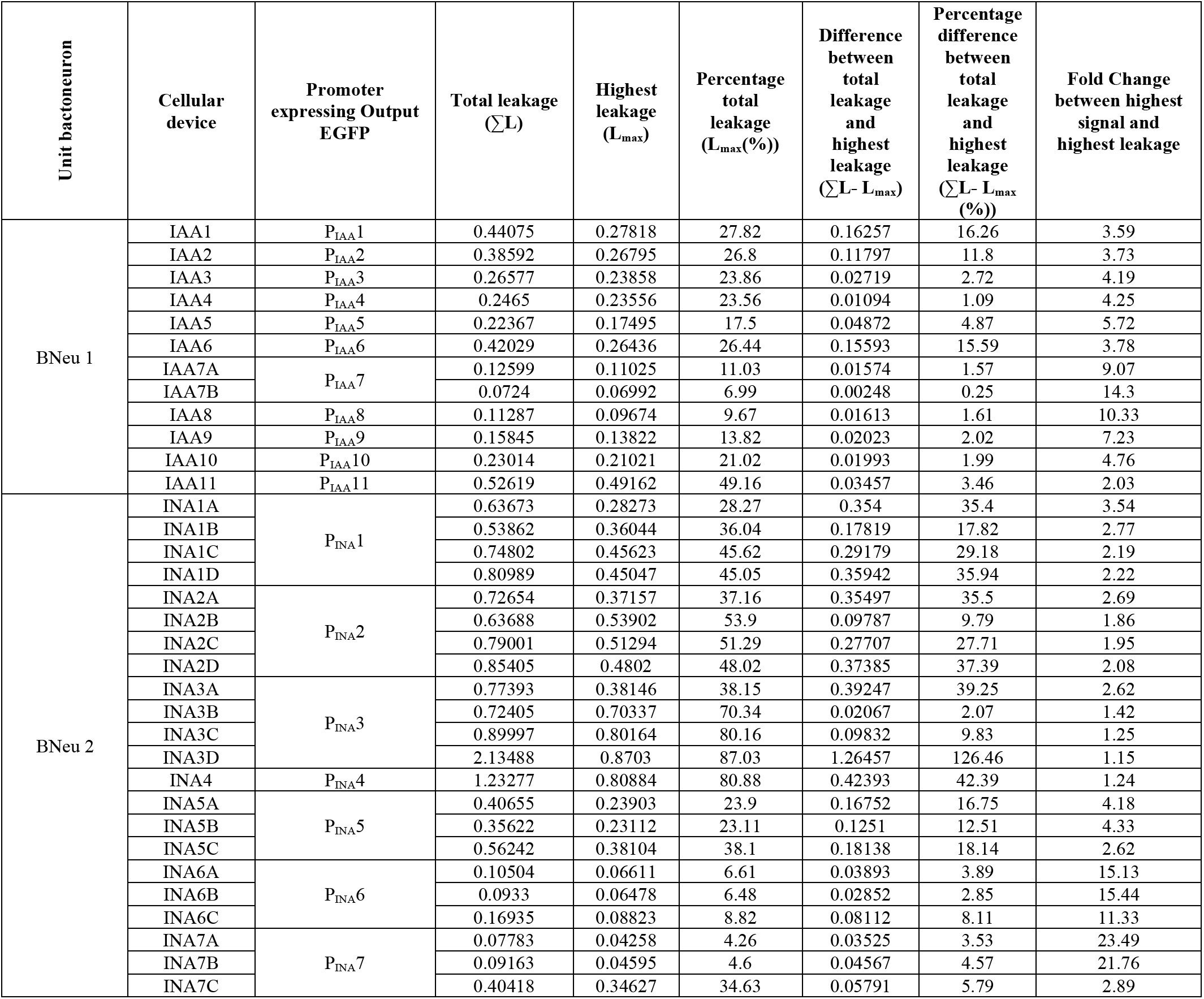
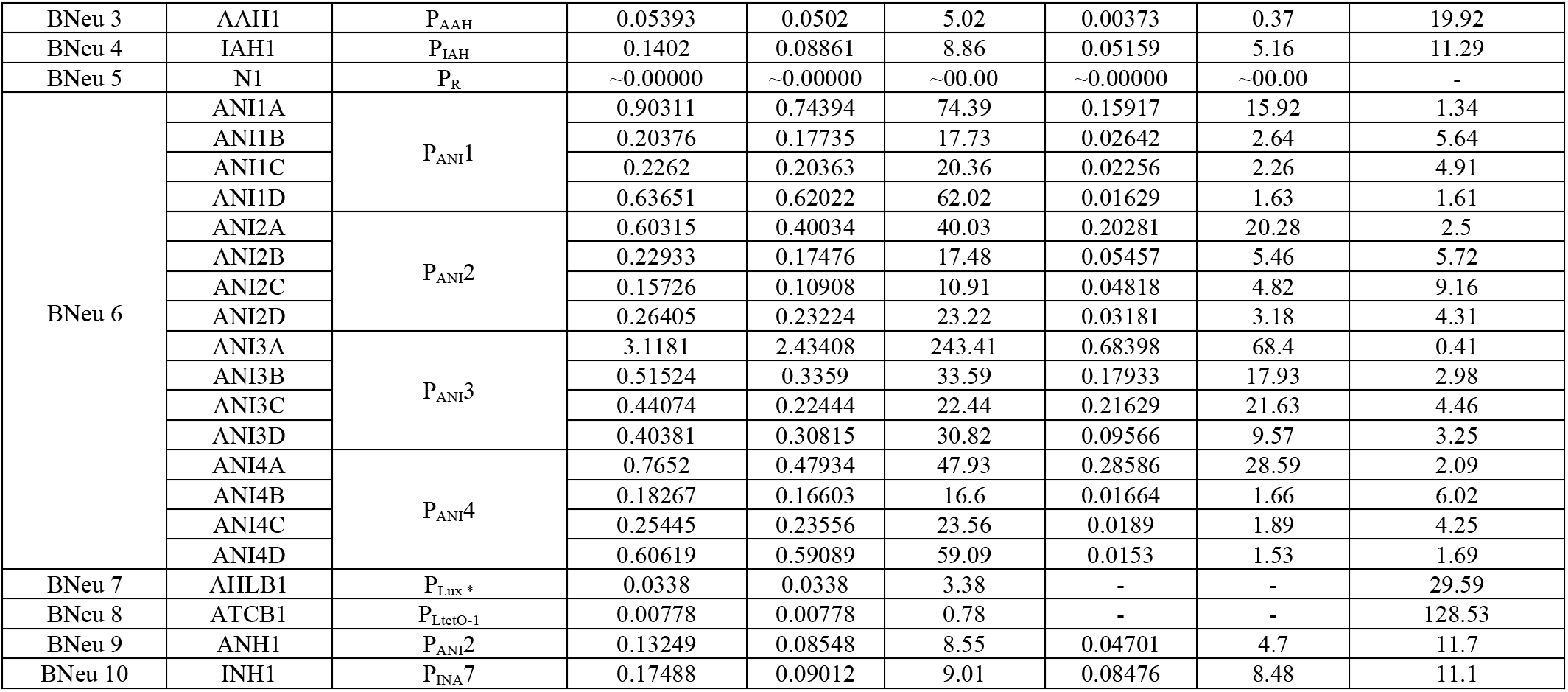
Leakage of each EGFP-expressing cellular device (construct) during weight and bias optimization of unit bactoneurons.

**Table S6:**
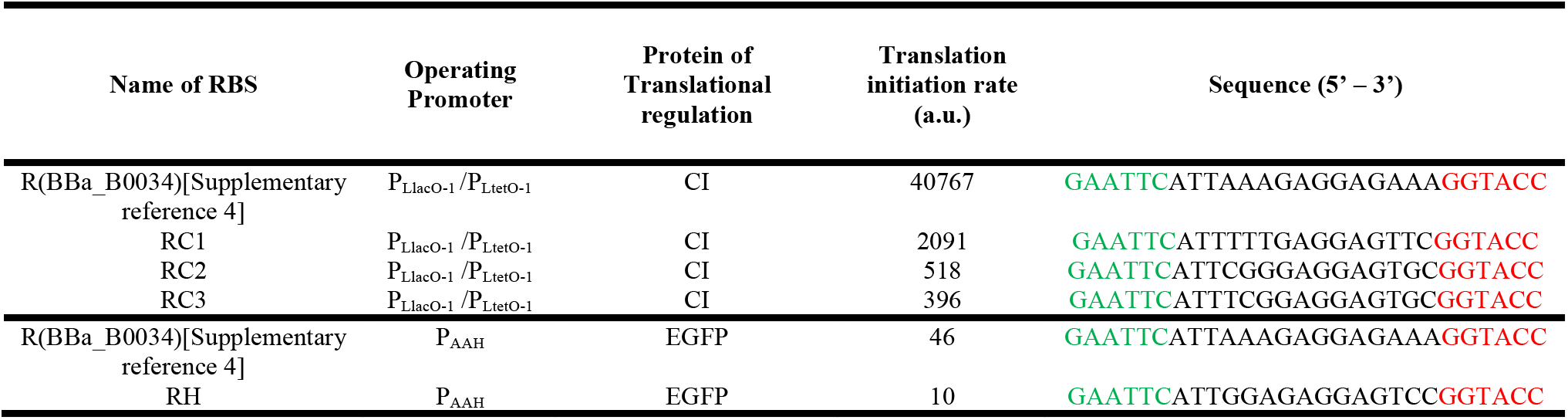
Translation initiation rate calculated from RBS calculator [Supplementary reference 3].

**Table S7:**
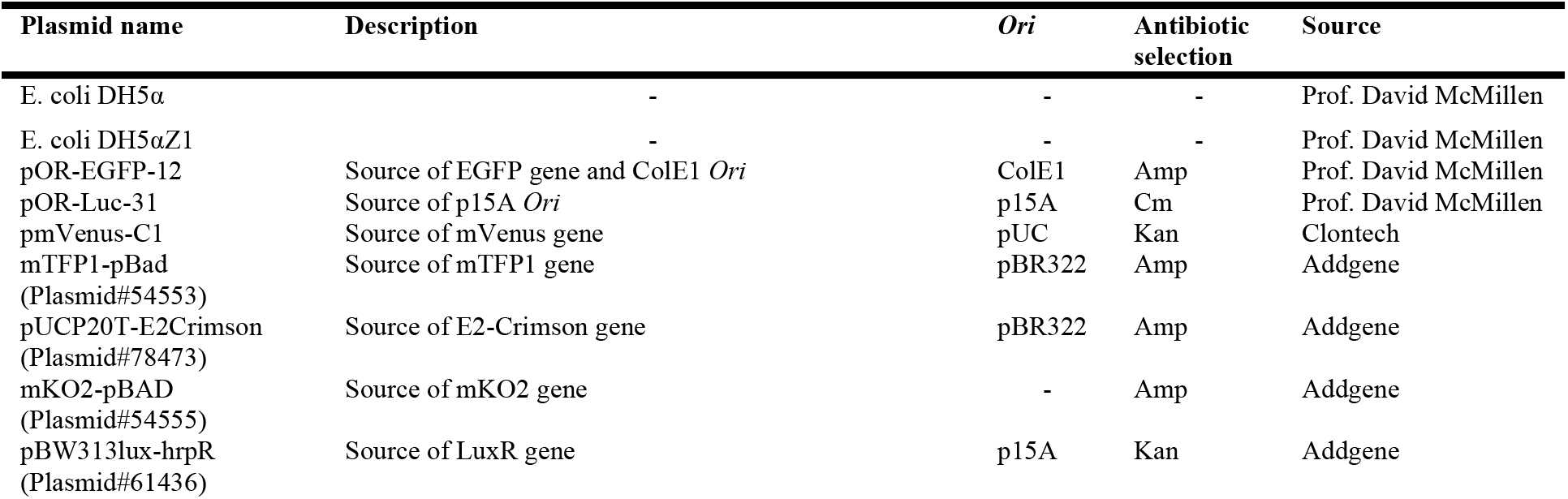
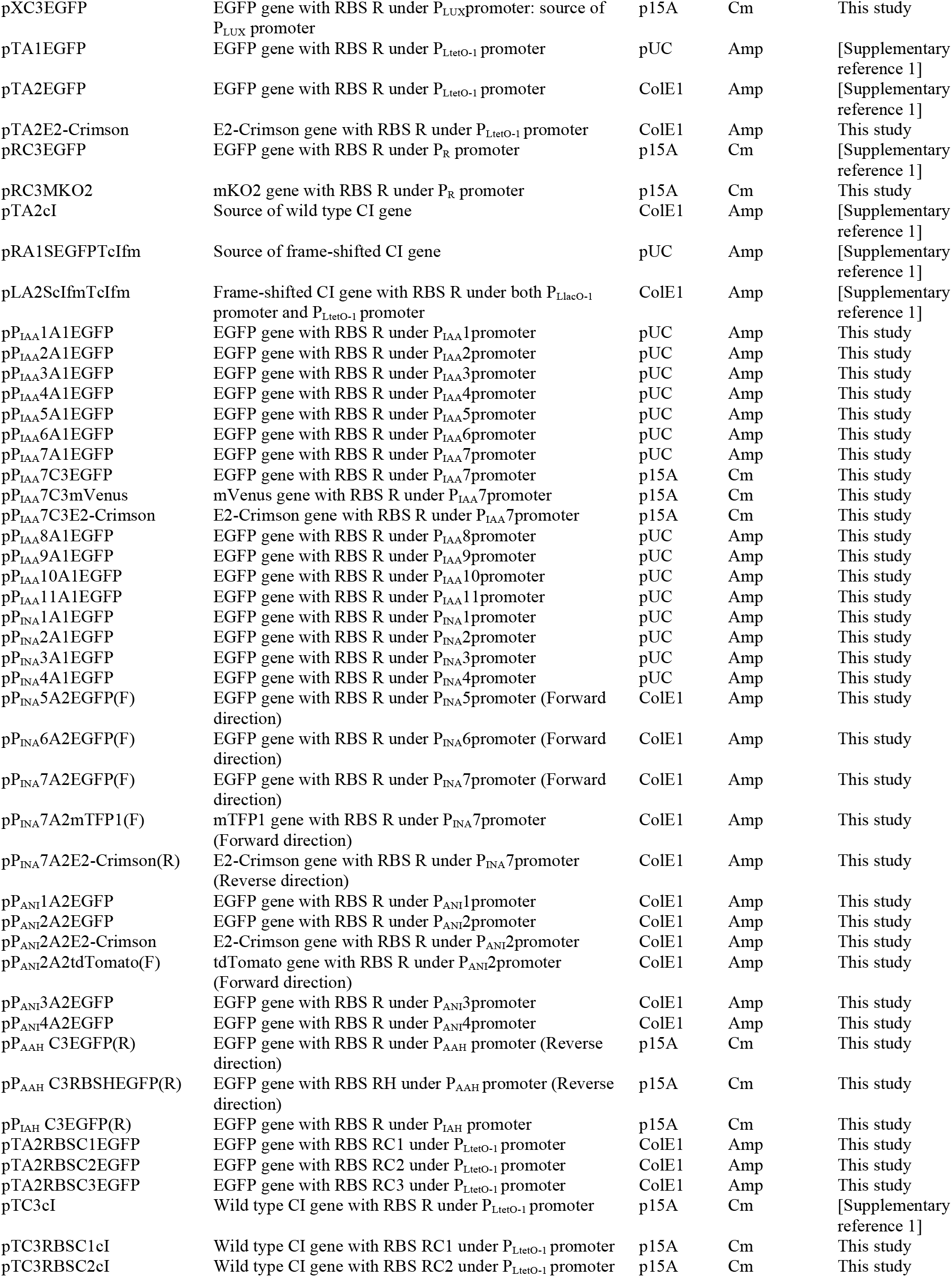
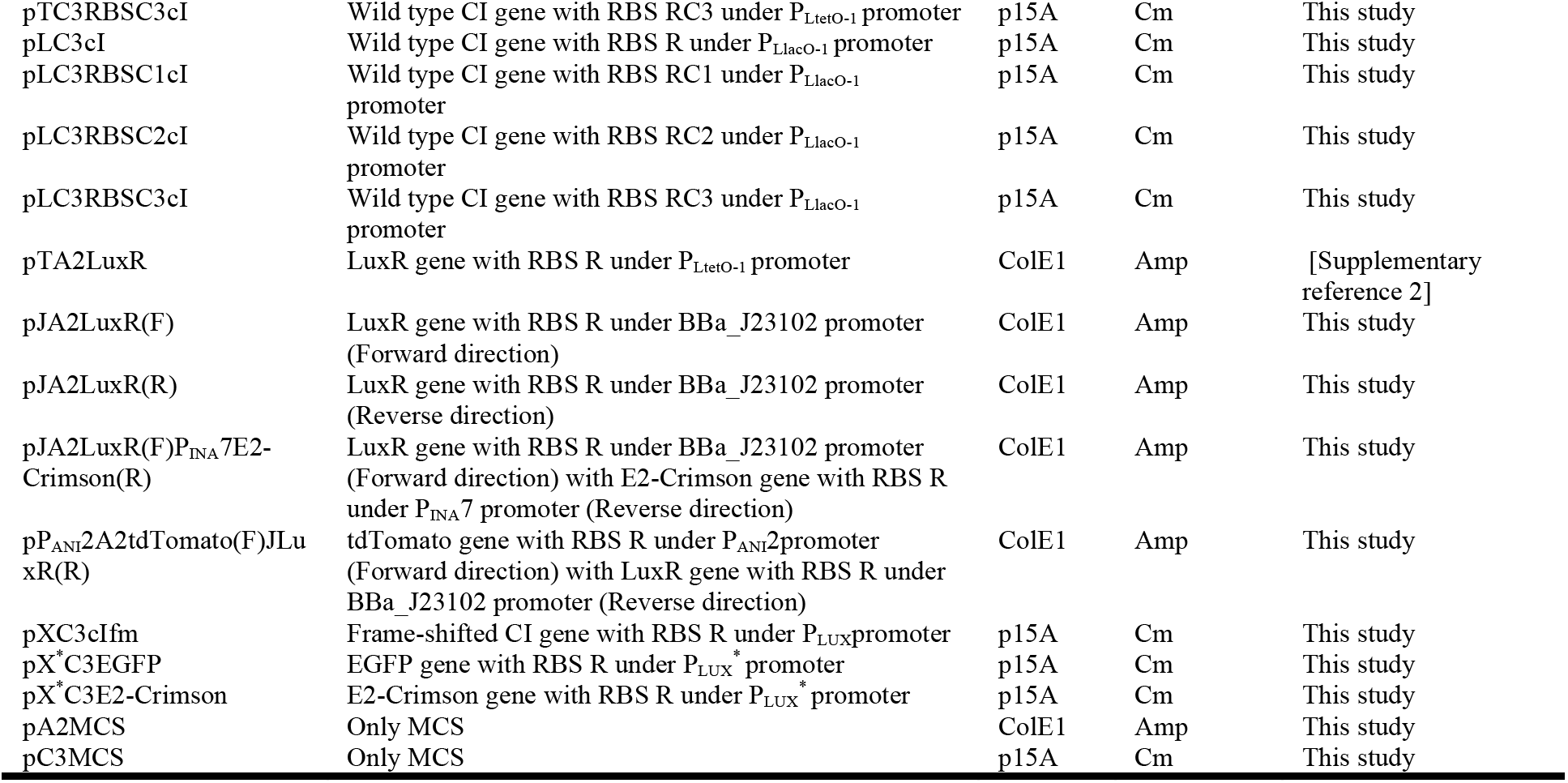
List of bacterial strains and plasmids used in this study.

## Notes

### Competing Interest Statement

The authors have declared no competing interest.

